# Fixed human pangenome sequences reveal origins of common human traits

**DOI:** 10.64898/2026.04.30.722098

**Authors:** Chern-Sing Goh, Matthew H. Davenport, Chul Lee, Erich D. Jarvis

**Author notes:** Corresponding: Chul Lee and Erich D. Jarvis. These authors contributed equally: Chern-Sing Goh, Matthew H. Davenport.

## Abstract

Humans possess human-specific traits^1,2^ such as spoken language and lineage-specific traits such as ape-specific taillessness^3,4^. Previous efforts to identify the DNA sequences responsible for such human traits were limited by necessary accommodations for poor genome assembly quality and lack of population genomic sampling^5–16^. Here, we implement new algorithms that combine the near-complete human reference pangenome alignment with a new near-complete simian cross-species alignment to define human- and lineage-specific DNA sequences fixed across human haplotypes. Previously reported *FOXP2/NOVA1*^17,18^ amino acid substitutions linked to human spoken language and *TBXT* transposable element insertion contributing to ape taillessness^3,4^ were unique to their respective clades and fixed in sampled humans. In contrast, widely used sets of candidate human-mutated loci showed limited enrichment for either human specificity or fixation. Integration with candidate *cis*-regulatory elements^19–22^ identified putative regulatory sequences specific to humans and linked to human-specific traits like hair reduction^23,24^ and brain transcriptome patterning^25^. Although brain-associated fixed regulatory changes were present in all lineages, enrichment for spoken language was human-specific and enrichment for receptive language was ape-specific. This study provides a new pangenome-aware comparative framework and catalogs of candidate genomic loci to trace the evolutionary origins of common human traits and disease risks.

## Main

Modern humans possess not only human-specific traits like spoken language, bipedalism, and reduced body hair^1,2^ but also lineage-specific traits like taillessness, which all apes share^3,4^. This evolutionary accumulation of human traits is thought to have involved trait-altering DNA changes, particularly in regulatory sequences, that arose along nested lineages and were subsequently retained or fixed in modern humans^26–28^. Therefore, a central challenge in evolutionary genomics is to identify modern human DNA sequences that are both lineage-specific and population-fixed, thereby most plausibly linked to common human traits^29,30^.

In the past decade, several comparative genomic studies have proposed candidate loci, most notably human accelerated regions (HARs)^5–10^ and human ancestor quickly evolved regions (HAQERs)^11–13^. However, the analytical tools used were developed in the context of short-read genome assemblies with lower accuracy and frequently misassembled regulatory sequences with high GC and/or repetitive content^14–16^. To reduce the impact of such artifacts, analyses often shifted from direct descriptions of individual mutations toward substitution-rate inferences within conserved or readily alignable genomic windows^5,6,8,11^. Although such approaches can nominate broad candidate regions^31,32^, they do not systematically identify the specific nucleotide changes fixed across modern humans within those regions. Further, previous comparative analyses relied on a single human reference genome, making them equally sensitive to private mutations in the reference individual as to common sequences shared by all humans^33^.

To address the underlying issues, sequencing consortia including the Vertebrate Genomes Project (VGP)^34,35^ and the Human Pangenome Reference Consortium (HPRC)^12,36^ have produced near-complete long-read genome assemblies of non-human simian species and geographically dispersed modern human individuals^37^. Here, we leverage this leap in data quality and coverage to identify human-fixed lineage-specific sequences (FLSS) at single-nucleotide resolution by developing a novel pangenome-aware comparative framework that integrates cross-species and within-species genome-wide analyses. The resulting FLSS catalogs, made available through a public UCSC track hub (see **Data availability**), reveal the evolutionary origins and genetic constraints of candidate *cis*-regulatory elements from the ENCODE project^19–22^ and brain-linked regulatory programs^25,38,39^, thereby connecting fixed human genome evolution to common human traits and neurocognitive disorders.

### Mosaic sequences invariant in humans

To trace the origins of human genome sequences that differ from those of other simians, we generated a 9-way cross-species alignment using near-complete genome assemblies from nine species spanning human to marmoset, capturing five nested lineages from human-specific to ape-wide branches, and paired it with a 464-way human pangenome alignment from 464 haplotypes of 232 globally sampled individuals with hg38 as a traditional reference to jointly assess lineage specificity and population invariance (**Extended Data Fig. 1a,b; Supplementary Tables S1-3; Methods**). To integrate these resources at base-pair resolution, we developed Genomic Origin tools (GOtools; https://github.com/csgDarwin/GOtools), a pangenome-aware comparative framework that uses hg38 as a common reference. GOtools identified lineage-specific sequences (LSS) based on perfect in-group retention and out-group absence at each base position in hg38, separately considering the in-groups and out-groups for each of the total five lineages, including humans (**Fig. 1a**). To assess selection, GOtools also identified all base positions in hg38 that were fully conserved across all haplotypes in the pangenome (autosomes: 464, chrX: 363, chrY: 118 haplotypes). Intersecting these sets yielded catalogs of FLSS of the five lineages (**Fig. 1b; Extended Data Fig. 2; Methods**).

**Figure 1.**
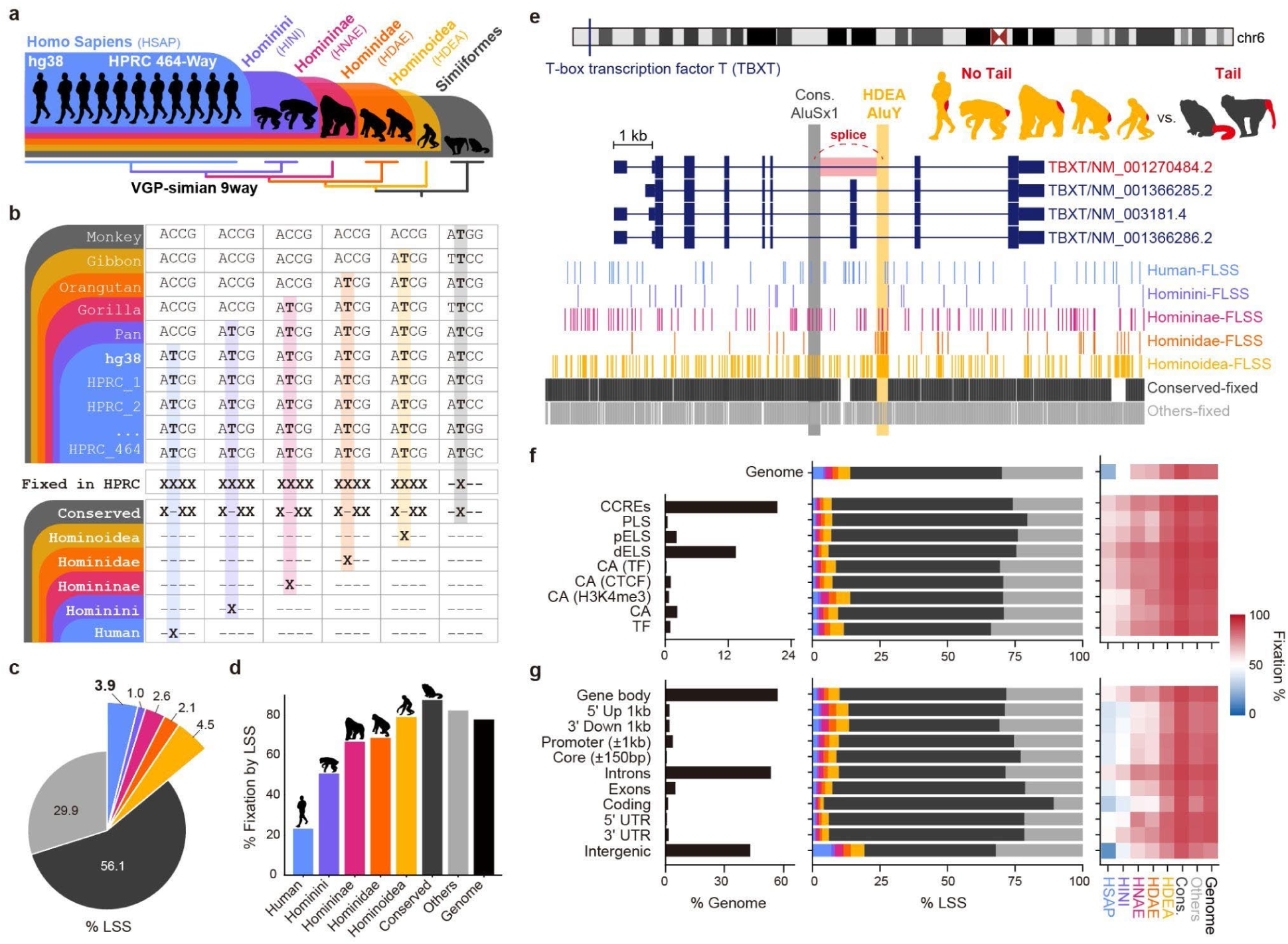
Identification of LSS and FLSS in hg38 reference. **a**, Schematic of the integrated HPRC 464-way human pangenome alignment and 9-way simian cross-species whole-genome alignment, both including hg38. **b**, Workflow for defining FLSS by intersecting human-fixed sequences from the HPRC 464-way alignment with human-specific or lineage-specific sequences (human-LSS, hominini-LSS, homininae-LSS, hominidae-LSS, and hominoidea-LSS) derived from the 9-way alignment. **c**, Genomic proportions of human-specific, lineage-specific, simian-conserved, and other sequences in the 9-way alignment. **d**, Proportions of each LSS fixed across the HPRC 464-way human pangenome. **e**, Example at *TBXT* showing a hominoidea-specific AluY insertion causing taillessness in apes. **f**, Genomic proportions of ENCODE cCRE categories, together with lineage-specific proportion (% LSS, stacked bars) and fixation rates (FLSS/LSS, heat map) for each lineage. **g**, Genomic proportions of genomic annotation categories, together with lineage-specific proportions (% LSS, stacked bars) and fixation rates (FLSS/LSS, heat map) for each LSS category. In **f** and **g**, the genome row summarizes the genome-wide values shown in **c** and **d**.

Human-specific sequences (human-LSS) and ape-specific sequences (hominoidea-LSS) comprised 3.9% and 4.5% of hg38, respectively, but differed markedly in pangenome fixation. Only 23% of human-LSS were fixed across sampled human haplotypes, compared with 79% of hominoidea-LSS, corresponding to 0.9% and 3.5% of hg38 being human-FLSS and hominoidea-FLSS, respectively. By contrast, most of the genome remained conserved across the 9-way simian alignment: 56.1% of hg38 was conserved, and 87.5% of that sequence was also fixed across the human pangenome (**Fig. 1c,d**; **Supplementary Table S4**).

### Fixation patterns in reported variants

Our FLSS analysis reidentified several mutations previously linked to lineage-restricted human traits. For language evolution, we found that the transcription factor *FOXP2* contains two single-nucleotide human-FLSS producing human-specific T303N and N325S amino acid substitutions^17^ and similarly found a coding single-nucleotide human-FLSS responsible for the known I197V substitution in the RNA-binding protein *NOVA1*^18^ (**Extended Data Fig. 3a,b**). In addition to these single-nucleotide mutations, our FLSS analysis reidentified the much larger AluY SINE insertion (290 bp) specific to apes in a 3’ intron of *TBXT,* which has been shown to contribute to tail loss in apes^4^ (**Fig. 1e**).

### Fixation rates in regulators and genes

We next measured the evolutionary rate by sequence category across human-simian ancestry by intersecting our FLSS sets with either ENCODE annotation of candidate cis-regulatory elements (cCREs)^19–22^ or RefSeq annotation-based genic structures^40^. The genome-wide pattern of older sequences with higher fixation rates held true across lineage categories, except in RefSeq-defined intergenic regions, which were enriched for human-LSS. However, these intergenic human-LSS positions in hg38 exhibited the lowest fixation rates of any sequence category by lineage combination in the pangenome (13.9%), which we expect due to an abundance of unselected intergenic transposons (**Fig. 1f-g; Supplementary Table S5**).

### HAR and HAQER contain a few human-FLSS

To benchmark against past approaches tracing human genome evolution, we next compared human-LSS and human-FLSS coverage across HARs^5–9,32^ (n = 3,257) and HAQERs^11^ (n = 2,747). Relative to the genome-wide background rates of 3.9% for human-LSS and 0.9% for human-FLSS, HARs were depleted for human-LSS (1.6%) and only slightly enriched for human-FLSS (1.3%). While HAQERs were enriched for both LSS and FLSS, the observed rates remained low (8.4% and 2.7%, respectively; **Extended Data Fig. 4a–d**). Examples from the longest accelerated regions further illustrate this distinction. The longest HAR, HAR_2635, showed sparse human-FLSS (0.5%), while the longest HAQER, T2T_HAQER_0215, did not contain any human-FLSS due to the absence of fixation in the human pangenome (**Extended Data Fig. 4e,f**).

This distinction was even more pronounced when examining the regulatory human-FLSS contained inside ENCDOE cCREs (**Fig. 2a**). Despite the relatively low human-unique sequence content observed in past models, >500-fold more cCREs (55.74% of cCREs) contained at least 1 bp of human-FLSS, compared with only 2,471 (0.11%) containing a HAR-fixed position and 1,585 (0.07%) containing a HAQER-fixed position (**Fig. 2b**). Similar to this difference in overlap, we also found that human-FLSS explained the evolution of 10-fold more fixed regulatory sequence (0.61% of cCRE bases) compared to HAR-fixed (0.06%) or HAQER-fixed (0.04%) positions (**Fig. 2c**). Thus, human-FLSS capture a substantially larger fraction of regulatory sequence evolution than similar HAR- or HAQER-sets. Within this context, the longest HAR-fixed cCRE (EH38E4058778, 341 bp) did not contain any human-LSS, whereas the longest HAQER-fixed cCRE (EH38E3253360, 317 bp) contained only 20 bp of human-LSS (6.3%) (**Extended Data Fig. 5a,b**).

**Figure 2.**
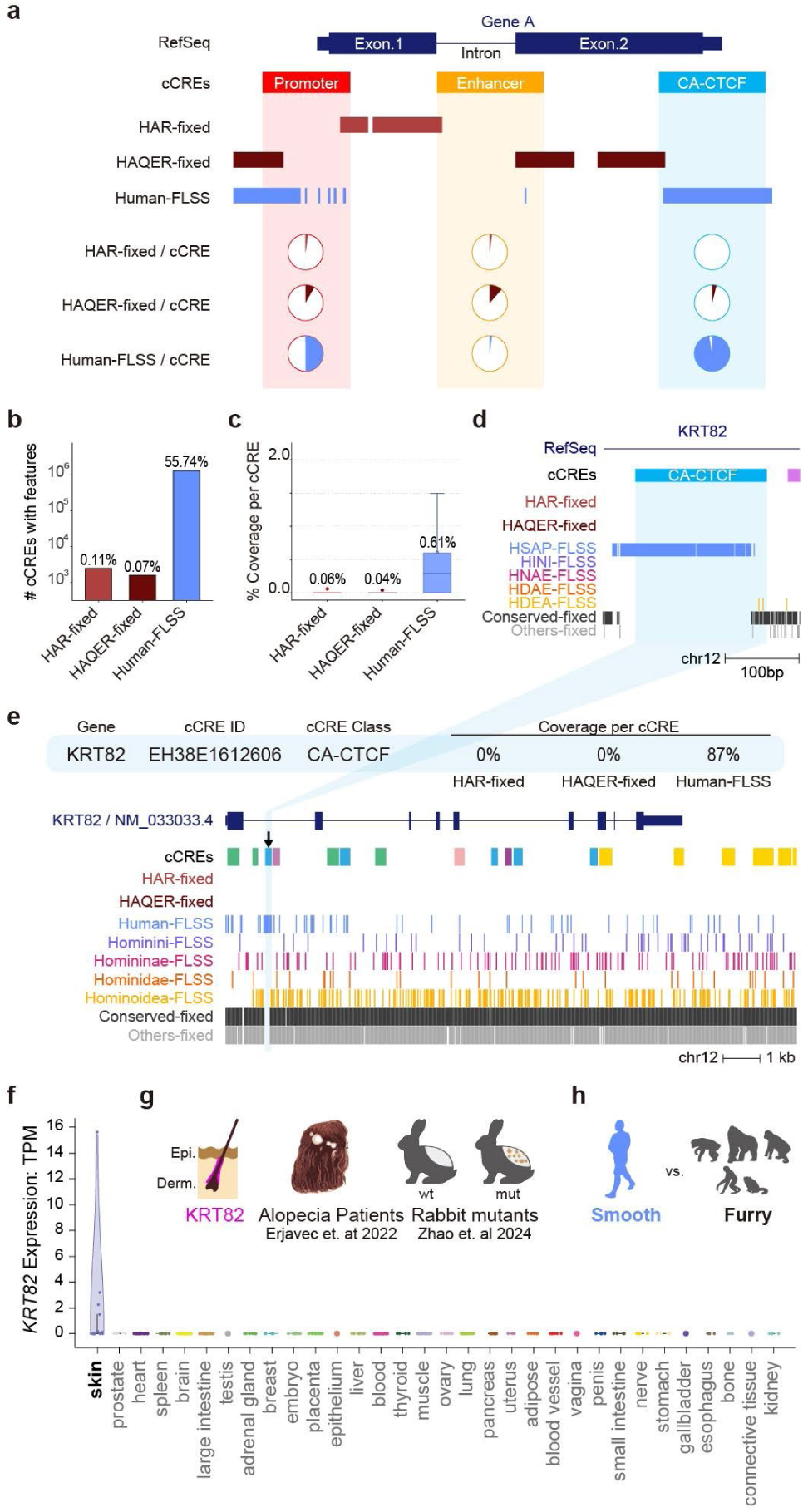
Integration of human-FLSS with ENCODE cCREs. **a**, Schematic showing the overlap of a hypothetical cCRE element with three fixed human variant datasets, HAR-fixed, HAQER-fixed, and human-FLSS. Pie charts indicate the fraction of positions within each cCRE that overlap each dataset. **b**, Number of hg38 cCREs containing at least one fixed variant. Values above bars indicate the number and percentage of cCREs that overlapped. **c**, Distribution of the percentage of positions per cCRE overlapping fixed variants. **d**, Genome browser view of an FLSS-rich CA-CTCF cCRE, EH38E1612606, in *KRT82.* **e**, Expanded genome browser view across the *KRT82* locus, showing the position of cCRE EH38E1612606 and the distribution of HAR-fixed, HAQER-fixed, and lineage-specific fixed variants. **f,** Expression summary for *KRT82* across human tissues in the SCREEN portal. **g**, Summary of published functional and expression contexts for *KRT82*. **h**, Schematic comparison of relative body hair coverage between humans and non-human simians.

### *De novo* human regulator for hairlessness

To prioritize human-specific and population-invariant cCREs linked to human-specific traits, we intersected with the ENCODE cCRE database to identify regulatory human-FLSS (**Supplementary Tables S5,S6**). As a simple example, we highlighted an FLSS-rich *CTCF*-binding element (87% human-FLSS in EH38E1612606) in an intron of *KRT82*, not previously identified by HARs or HAQERs (**Fig. 2d,e**). This *CTCF*-binding element was created by a 176 bp human-specific insertion absent in other non-human simians. As confirmation, BLAT mapping of the flanking human sequences to the chimpanzee assembly was consistent with the human-specific insertion (**Extended Data Fig. 6a**). *KRT82* is a hair keratin expressed by hair shaft cells in hair follicles during hair growth^23^ and whose expression is highly specific to skin at the tissue level (**Fig. 2f**). *KRT82* was also the top candidate from a recent GWAS study into the genetic basis of human alopecia^23^, and a *KRT82* promoter allele in rabbits produces a “patchy fur” phenotype^24^ (**Fig. 2g**). These observations of *KRT82* expression affecting hair patterning prioritize a further validation test for the human-specific and fixed *CTCF*-binding element in different hair regulations on human scalp and body (**Fig. 2h**).

### Drivers changing human brain expression

As the phenotypes most often thought of as uniquely human are behavioral (language, cognition, tool use, etc.) and thus brain-derived, we integrated our % regulatory human-FLSS dataset with publicly available comparative brain single-nucleus RNA sequencing (snRNAseq) data. Specifically, we compared RNA abundance from homologous brain tissue (middle temporal gyrus) in humans, chimpanzees, and gorillas^41^ to identify human-altered expression at the cell-type level (**Fig. 3a**; see **Methods**). This analysis identified 10,232 genes differentially expressed in at least one cell type in the human brain, with the Layer2/3 intratelencephalic glutamatergic (IT-glut) neuron subtype being the most altered cell type in these data (**Fig. 3a**). To find patterns in these data, we built a matrix of human differential expression direction by cell type and performed hierarchical clustering to identify gene sets with consistent differential expression patterns across cell types (**Fig. 3b, Supplementary Table S7**). For example, one set of genes was upregulated in human IT types across layers but unaltered in glia, whereas another set was upregulated only in human astrocytes. To build human evolutionary models of the differential expression of these genes, we then sorted the % human-FLSS cCRE data for each differentially patterned gene set to identify human-FLSS rich regulatory regions (**Fig. 3c**).

**Figure 3.**
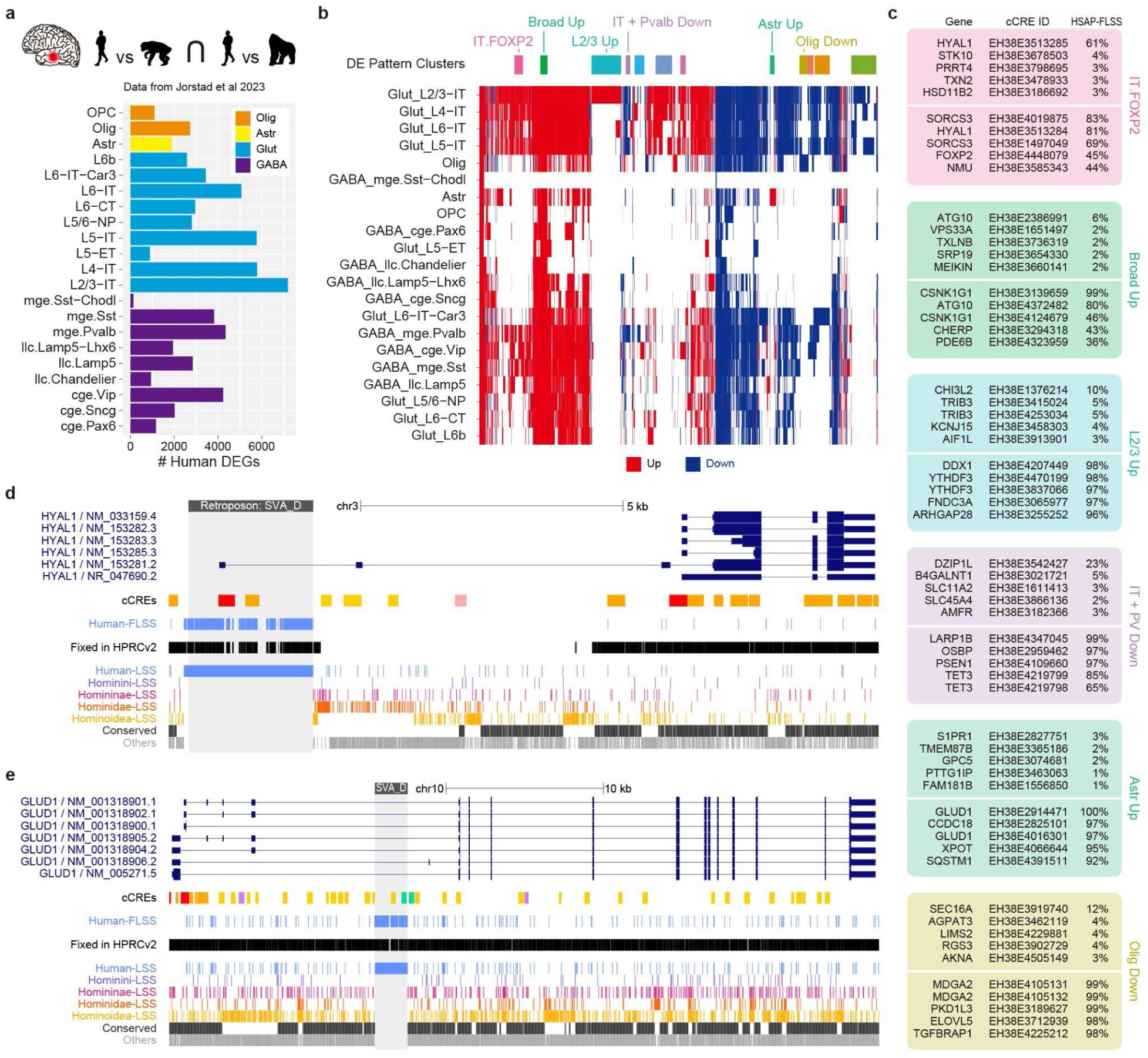
Regulatory FLSS altered human brain gene expression. **a**, Schematic of differential testing between human, chimpanzee, and gorilla brain cell types dissected from the midtemporal gyrus and the number of human differentially expressed genes (DEGs) per cell type. **b**, Hierarchical clustered matrix of DEGs to identify genes with similar patterns of differential expression across cell types. **c**, cCREs for each differential expression pattern cluster ranked by percentage of human-FLSS overlap. Promoters (tops) and enhancers (bottoms) were considered separately. **d**, Locus view of *HYAL1* showing transcript models, ENCODE cCRE annotations, human-FLSS, LSS, fixed sequence, conserved, and other sequence tracks. **e**, Locus view of *GLUD1* showing the same data tracks as **d**.

Among the genes upregulated in human IT neurons, *HYAL1* emerged as a notable candidate, with a promoter containing 61% human-FLSS and an enhancer containing 81% human-FLSS, both inserted into human *HYAL1* by an SVA_D retroposon (**Fig. 3c,d**). This new promoter created a long human-specific transcript with multiple newly expressed exons (**Fig. 3d**, NM_153281.2). BLAT mapping of the flanking sequence is consistent with the absence of this insertion in the chimpanzee genome (**Extended Data Fig. 6b**). Cross-species browser views further place this locus in a human-upregulated context in IT neurons (**Extended Data Fig. 7**).

From the genes upregulated in human astrocytes, the neurotransmitter glutamate-degrading enzyme *GLUD1* contained two human-FLSS-rich enhancers (100% and 97%) that were inserted together into human *GLUD1* by another SVA_D retroposon (**Fig. 3c,e**). Cross-species snRNA-seq data show that *GLUD1* is highly expressed in astrocytes across all simians, with even higher expression in humans (**Extended Data Fig. 8**). BLAT mapping is consistent with the absence of the corresponding insertion in the chimpanzee assembly (**Extended Data Fig. 6c**). These examples show that human-FLSS-rich regulatory elements are associated with human-specific brain gene expressions.

### Group functions of regulatory FLSS genes

To assess the group function of the genes most influenced by regulatory FLSS, we next performed gene set enrichment analysis on the top 1,000 genes ranked by the number of regulatory FLSS bases in each lineage (**Fig. 4a; Supplementary Tables S8-14; Methods**). Neuronal expression was enriched in the regulatory FLSS of each lineage with cellular compartment terms such as neuron projection, synapse, and dendrite (**Fig. 4b**). Enrichments in the KEGG database of molecular pathways better revealed lineage structure, with dopaminergic signaling enriched only in humans. Several pathways, including axon guidance and morphine addiction, were enriched across all regulatory FLSS sets except for great apes (hominidae; HDAE). Others, including glutamatergic synapse, calcium signaling, long-term potentiation, and long-term depression, were enriched in some combination of top regulatory FLSS genes of humans, African great apes (homininae; HNAE), and apes (hominoidea; HDEA) (**Fig. 4c**).

**Figure 4.**
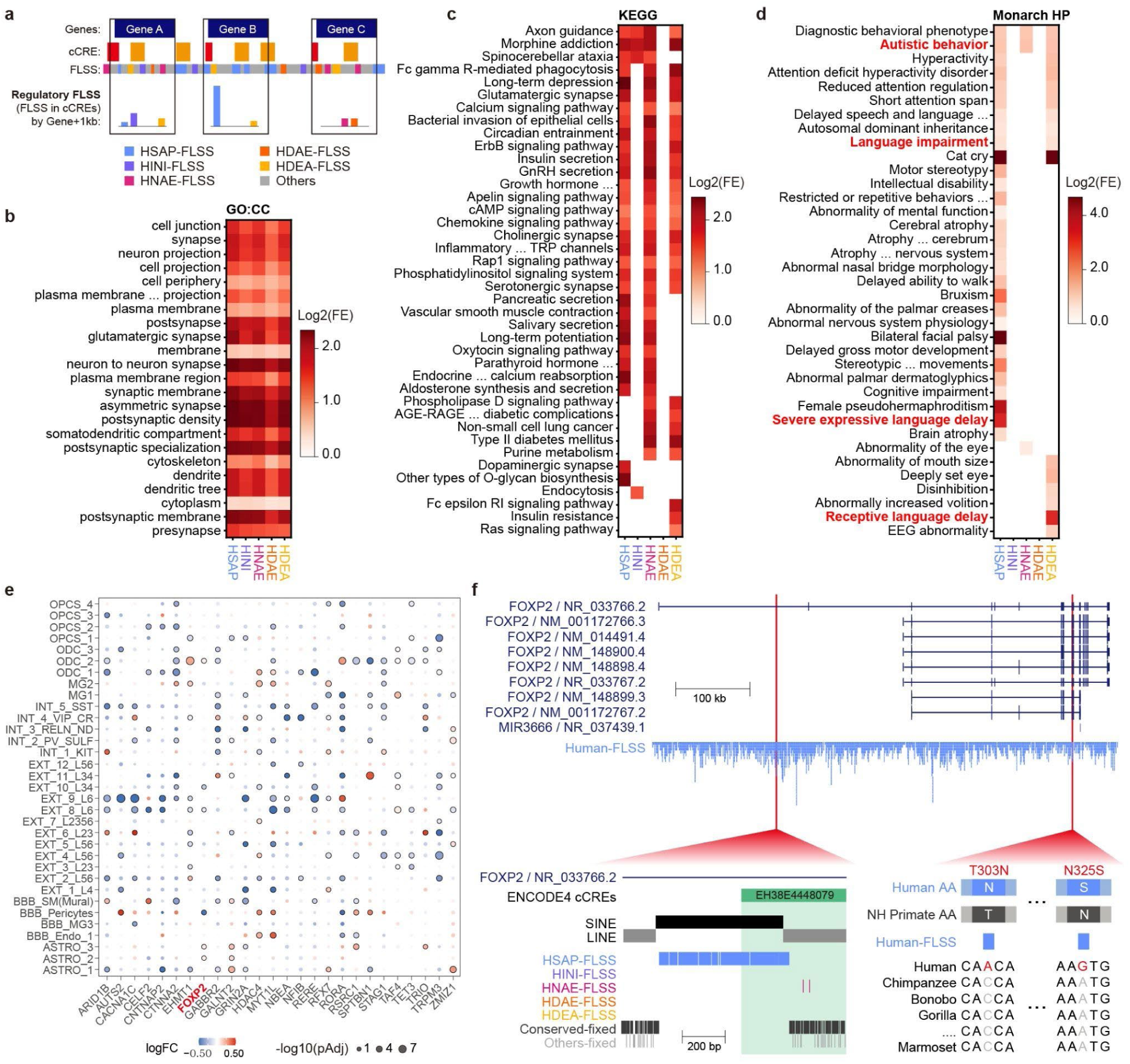
Top regulatory FLSS genes are linked to human phenotypic abnormalities. **a**, Schematic of the framework used to rank genes according to regulatory FLSS content within gene bodies and 1kb upstream promoters. Bar charts show the regulatory FLSS counts by lineage for three hypothetical genes. **b**, Heatmap of Gene Ontology Cellular Component (GO:CC) term enrichments for the top 1,000 genes based on FLSS content for each lineage. **c**, Heatmap of KEGG pathway enrichment for the same ranked gene sets across lineages. **d**, Heatmap of Human Phenotype Ontology (HP) enrichment= for the same ranked gene sets across lineages. All enrichments were log2 transformed for visual clarity. **e**, Dot plot describing altered expression in postmortem ASD patient brains (prefrontal cortex) for a high confidence set derived by the intersection of earlier analyses. **f**, Locus view of *FOXP2* showing transcript models, ENCODE cCREs, and FLSS tracks highlighting an FLSS-rich intronic enhancer, EH38E4448079, and two coding mutations.

### Human disease risks from lineages

We next used regulatory FLSS to better understand the evolution of human genetic disease by examining enrichments in the Human Phenotype Ontology database of clinical phenotypes and associated genetic variation. This analysis found that most clinical phenotypes aligned with regulatory FLSS emerged in either human or ape lineages, while fewer significant enrichments were observed in African great apes, and none in other lineages (**Fig. 4d**). Broadly, these enrichments describe fixed genomic signatures for the evolution of human hand and face morphology, and motor (e.g., walking), cognitive, and vocal behaviors. Human phenotype enrichments for regulatory human-FLSS included language impairment, neurodegeneration, cognitive impairment, motor stereotypy, developmental delay, and craniofacial abnormalities. Broad autistic behavior and diagnostic behavioral phenotype terms were enriched among the top regulatory FLSS genes in humans, African great apes, and apes. Several attention or language-related terms were shared between humans and apes. Interestingly, receptive language delay (language comprehension) was exclusively enriched in apes, whereas severe expressive language delay (spoken language) was enriched only in humans.

### Human regulatory evolution in autism

To produce a small high-confidence set related to human regulatory evolution and ASD, we intersected gene sets from enriched human phenotype terms of language impairment/autistic behavior and SFARI^42^ ASD-risk with our earlier analysis of differentially expressed genes in the human brain compared to apes (**Supplementary Table S15**). This analysis highlighted 26 genes that were highly affected in brain snRNAseq from ASD patients, using ASD-snGENE portal^43^ (**Fig. 4e**). STRING^44^ analysis showed that this gene set, including *FOXP2* and *CNTNAP2*, forms an interaction network involved in brain development, consistent with shared neurodevelopmental disease risk (**Extended Data Fig. 9**).

### Human evolution in *FOXP2*

A representative example from this prioritized set is *FOXP2*. In addition to the previously described human-FLSS amino acid substitutions (**Fig. 4f, Extended Data Fig. 3a**), the gene contains a previously undescribed intronic enhancer with 45% regulatory human-FLSS (EH38E4448079) identified by IT-upregulated differential expression analysis (**Fig. 3c**) corresponding to a human-specific SINE insertion (**Fig. 4f; Extended Data Fig. 6d**). Cross-species browser views further place this region in a human-upregulated IT-neuron expression context (**Extended Data Fig. 10**).

## Discussion

By integrating a cross-species simian genome alignment with the human pangenome, we establish a framework for interpreting the human genome as a mosaic of sequences from various phylogenetic origins, with fixation across modern humans. We achieved this by developing a simple-but-strict algorithm to reveal the lineage specificity and population commonality of human DNA sequences at single-nucleotide resolution. This framework required each nucleotide to be present in all extant species of a given simian lineage, including all globally sampled human pangenome haplotypes, and absent in each corresponding sister taxa. This analysis was made possible by the improvements in genome assembly quality achieved over the past decade through long-read sequencing technology. Applying this framework to a lower-quality alignment of short-read genome assemblies would be difficult, because biological variation could be confounded by missing GC-rich promoters and other technical artifacts in complex regulatory regions. Window-based approaches such as HAR and HAQER analyses were developed to mitigate these limitations, but contemporary genomic resources now enable nucleotide-resolved human comparative analyses like our pangenome-aware FLSS identification.

Beyond the gene-level examples described here, including *FOXP2, NOVA1, TBXT1, KRT82, HYAL1,* and *GLUD1*, we provide access to the full nucleotide-resolved FLSS catalogs through a UCSC track hub for Human Genome Sequence Evolution (HGSE; see **Data availability**). This public track hub allows users to integrate FLSS annotations with external datasets and easily generate publication-quality graphics describing any gene of interest. We also implement this evidentiary standard, in-group fixation and out-group absence, in GOtools for species with suitable cross-species and pangenome alignments of near-complete assemblies.

Through the progress made here, we identified several knowledge gaps to be addressed in future work. We used hg38 as the reference due to the abundance of associated resources (e.g., ENCODE cCREs), but future analyses using a telomere-to-telomere assembly such as hs1 would extend FLSS discovery into difficult-to-assemble regions, including telomeres and centromeres. The lack of HPRC-quality pangenome resources for non-human apes, especially chimpanzees, limits our ability to distinguish unsampled variations within populations of closely related species. The human-referenced FLSS catalogs contain lineage-specific substitutions and insertions, but not deletions^45,46^. As the genome alignments used here were not codon-aware, amino acid variants would be more accurately detected using gene alignments of protein-coding sequences to avoid alignment errors such as false frameshifts. Most importantly, all inferences presented here remain correlative. Extensive experimental work in genome-edited human cells and organoids *in vitro,* and appropriate *in vivo* models will be required to reasonably determine whether and how any given FLSS contributes to candidate human traits.

Building on these results, we propose that long-read genomics will shift evolutionary analyses away from broad candidate regions toward nucleotide-resolved mutations whose functional effects can be individually modeled and experimentally validated. Such efforts may inform genome-guided approaches to human biology and precision medicine, but they first require accurate descriptions of derived sequence states^47^, such as those provided here for human evolution.

## Extended Data Figures

**Extended Data Figure 1.**
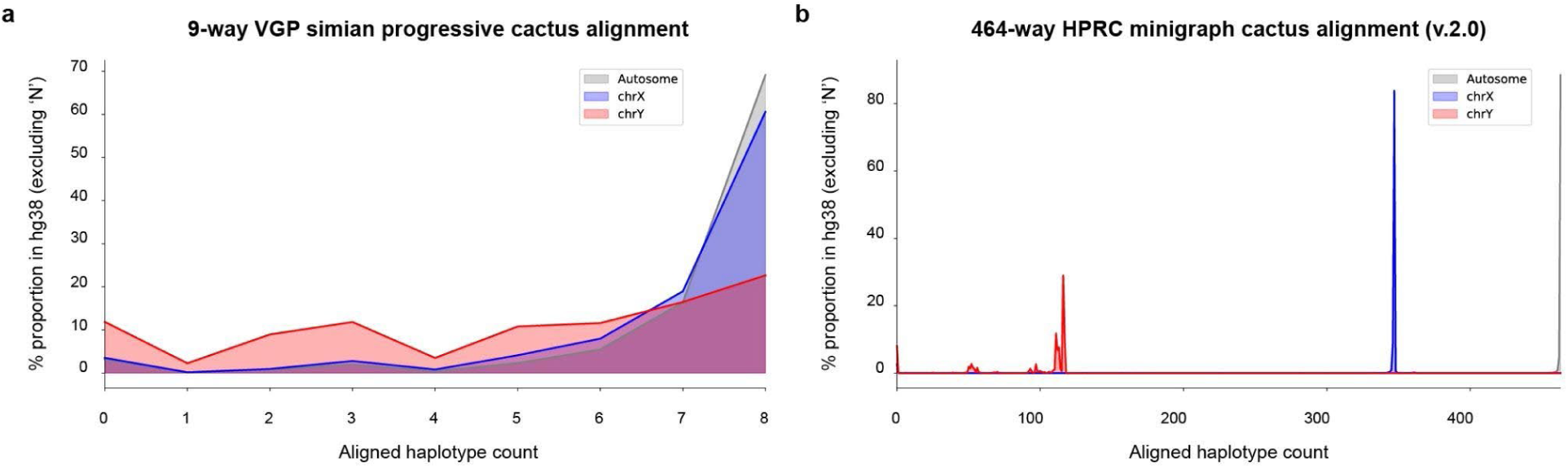
Distribution of aligned haplotype counts in both alignments. **a**, Distribution of aligned haplotype counts in the 9-way VGP simian progressive cactus alignment for autosome, X-chromosome, and Y-chromosome sequences, shown as the percentage of hg38 bases excluding Ns. **b**, Distribution of aligned haplotype counts in the 464-way HPRC minigraph-cactus alignment (v2.0) for autosomal (grey), X-chromosomal (blue), and Y-chromosomal (red) sequences, shown as the percentage of hg38 bases excluding Ns. The peak positions for sex chromosomes reflect their lower maximum aligned haplotype counts relative to autosomes (chrX: 363; chrY: 118; autosomes: 464).

**Extended Data Figure 2.**
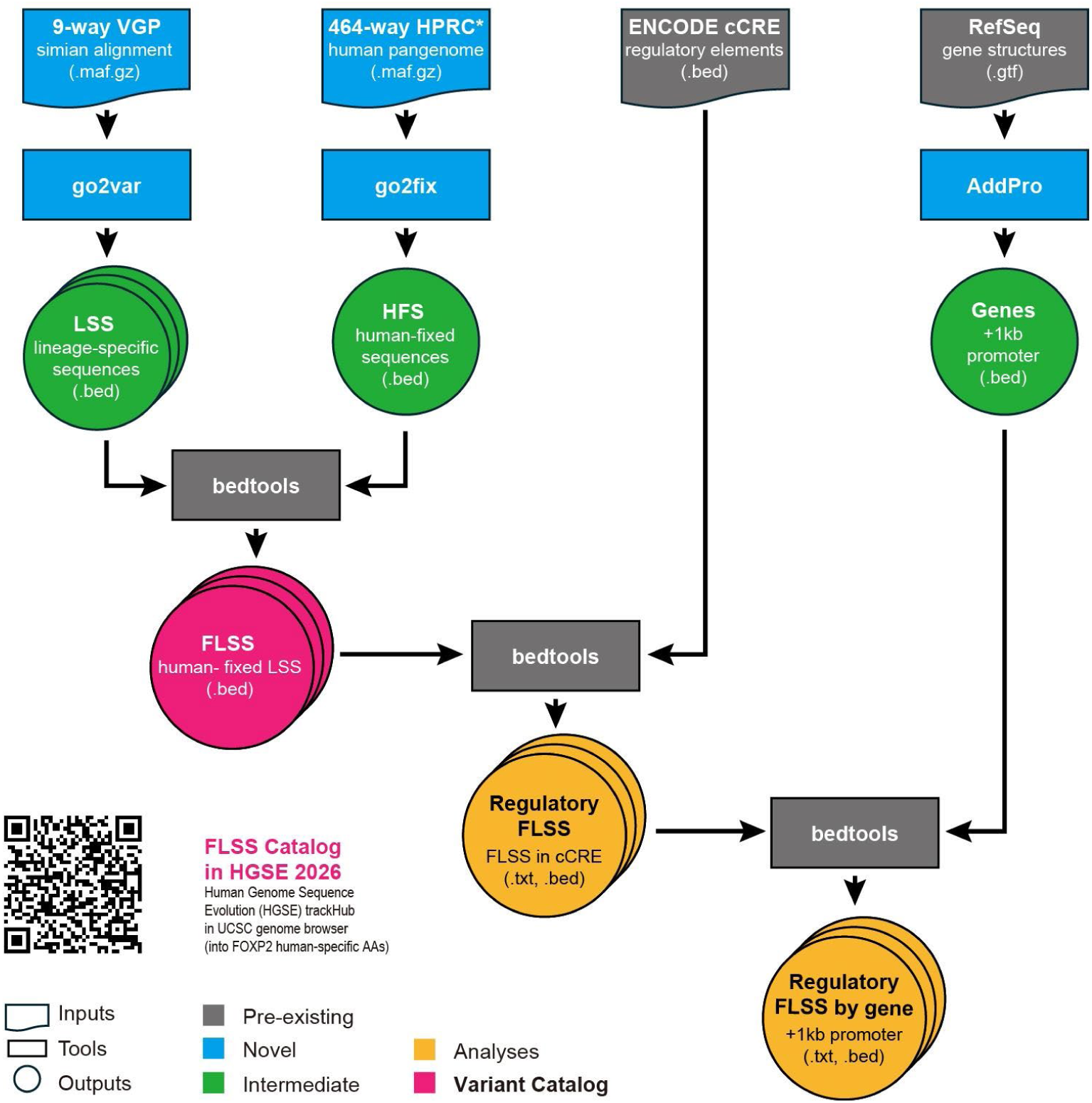
Schematic of computational pipeline. The 9-way simian haploid alignment is processed with go2var to generate lineage-specific sequences (LSS), and the 464-way human pangenome diploid alignment (HPRC v2.0) is processed with go2fix to generate human-fixed sequences (HFS). ENCODE cCREs were downloaded from the SCREEN database, and RefSeq gene annotations are extended to include 1-kb upstream promoter regions using AddPro. The intersection of HFS and LSS defines fixed lineage-specific sequences (FLSS). FLSS positions are then intersected with cCREs to generate regulatory FLSS, and subsequently with gene regions to generate the final per-gene output, regulatory FLSS by gene.

**Extended Data Figure 3.**
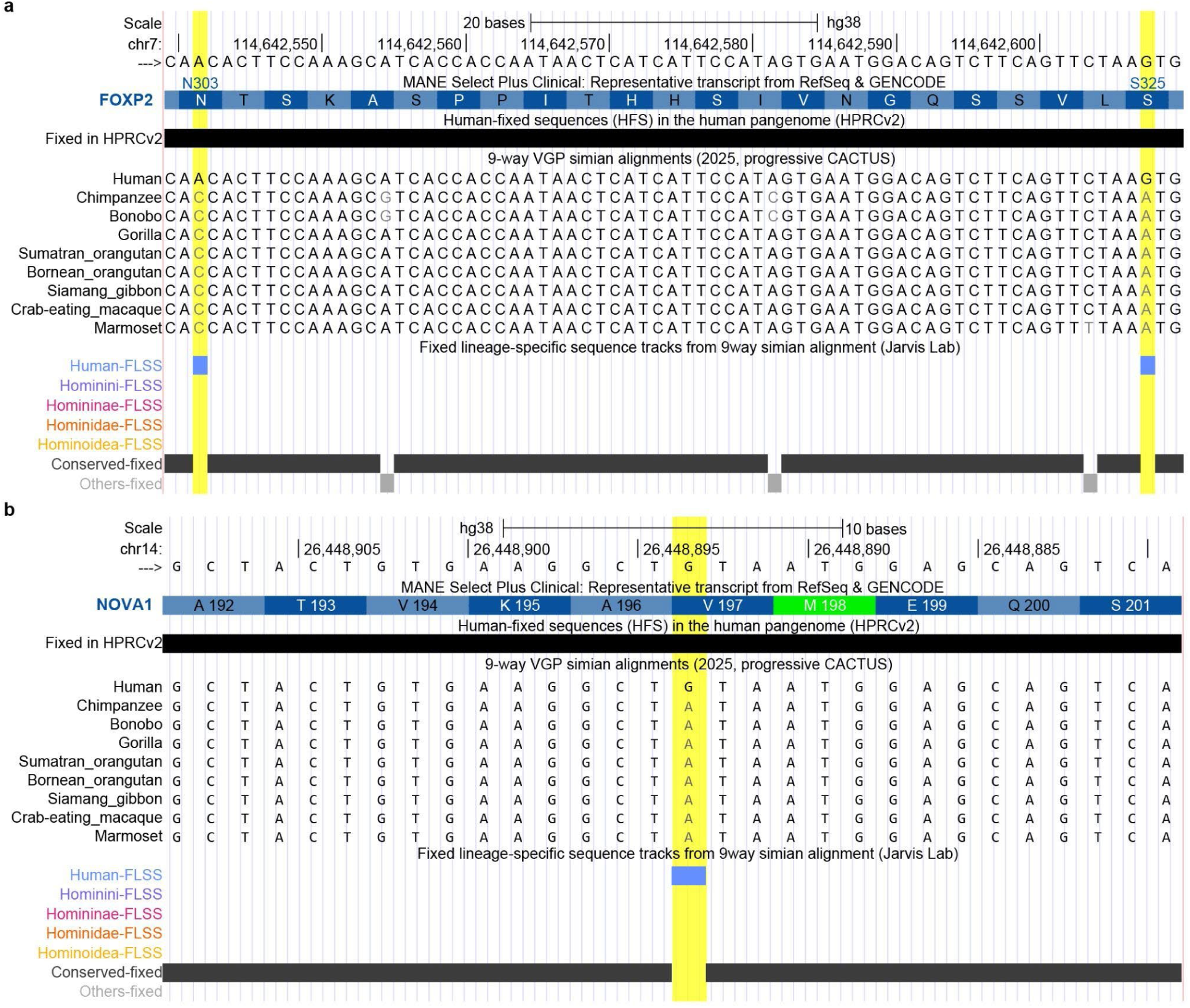
Fixation of human-specific missense variants in *FOXP2* and *NOVA1* in the human pangenome. **a**, UCSC Genome Browser view of the *FOXP2* coding region on chr7:114,642,540–114,642,610 (hg38), spanning amino acids 303–325 and containing the human-specific missense substitutions T303N and N325S. Both mutations were identified by GOtools as human-FLSS. **b**, UCSC Genome Browser view of the *NOVA1* coding region on chr14:26,448,880–26,448,909 (hg38), centered on amino acid 197 and the human-specific missense variant I197V. This site was identified by GOtools as human-FLSS.

**Extended Data Figure 4.**
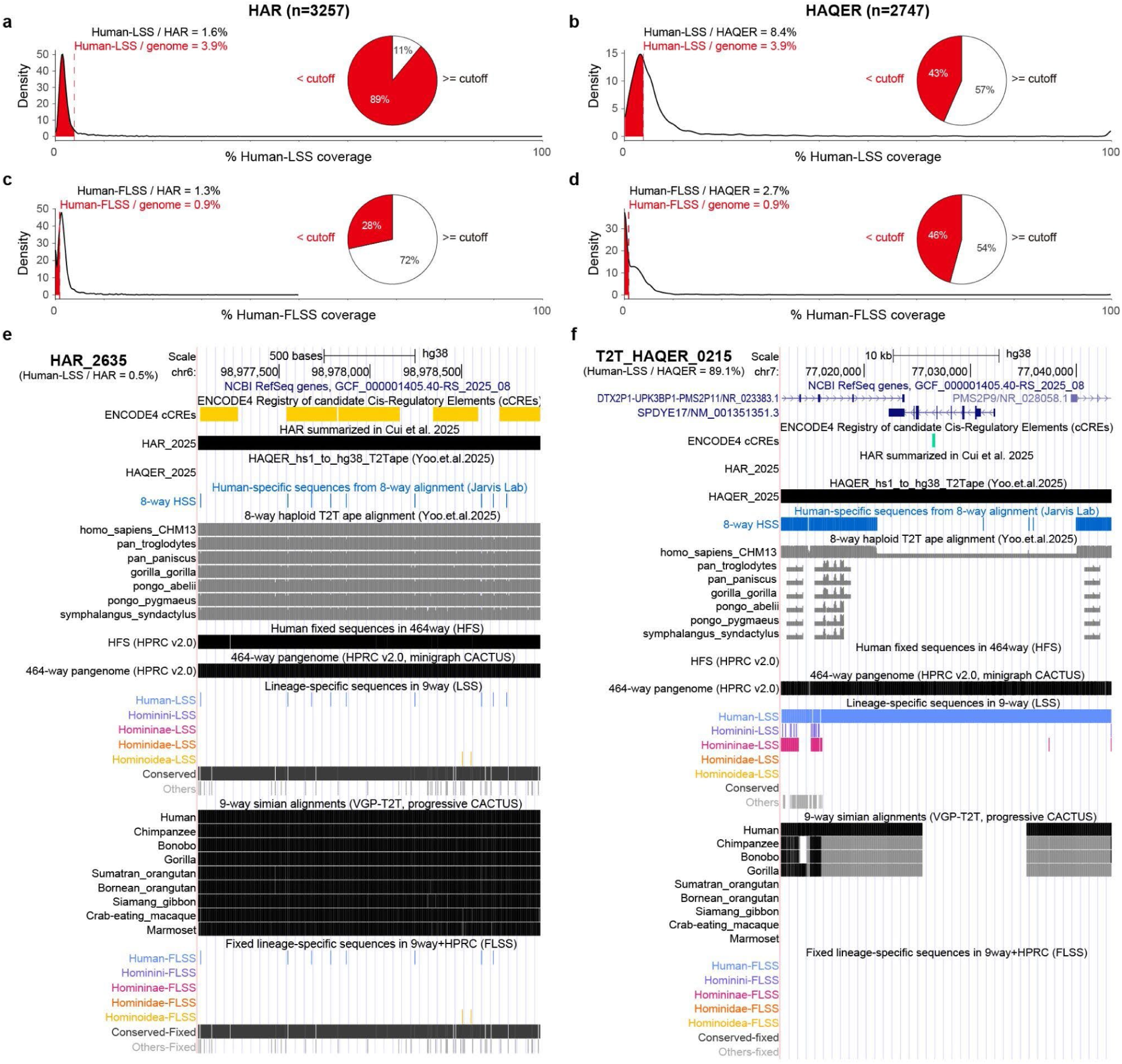
Human-LSS and human-FLSS coverage across HARs and HAQERs. **a**, Distribution of human-LSS coverage across Human Accelerated Regions (HARs; n = 3,257), measured as the percentage of bases in each HAR overlapping human-LSS positions. Black and red labels indicate the mean human-LSS coverage across HARs and the genome-wide background, respectively. The inset describes the HARs below the genome-wide background. **b**, Distribution of human-LSS coverage across Human Ancestor Quickly Evolved Regions (HAQERs; n = 2,747), labeled as in **a**. **c**, Distribution of human-FLSS coverage across HARs (n = 3,257), measured as the percentage of bases in each HAR overlapping fixed human-specific positions. Black and red labels indicate the mean human-FLSS coverage across HARs and the genome-wide background, respectively. The inset describes the HARs below the genome-wide background. **d**, Distribution of human-FLSS coverage across HAQERs (n = 2,747), labeled as in **c**. **e**, UCSC Genome Browser view of HAR_2635 (chr6:98,977,500–98,978,500; hg38), the longest HAR in the set. This region shows 0.5% human-LSS coverage and sparse human-FLSS positions. **f**, UCSC Genome Browser view of T2T_HAQER_0215 (chr7:77,020,000–77,040,000; hg38), the longest HAQER in the set. This region shows 89.1% human-LSS coverage but no human-FLSS, consistent with the absence of human-fixed sequence in the HFS track.

**Extended Data Figure 5.**
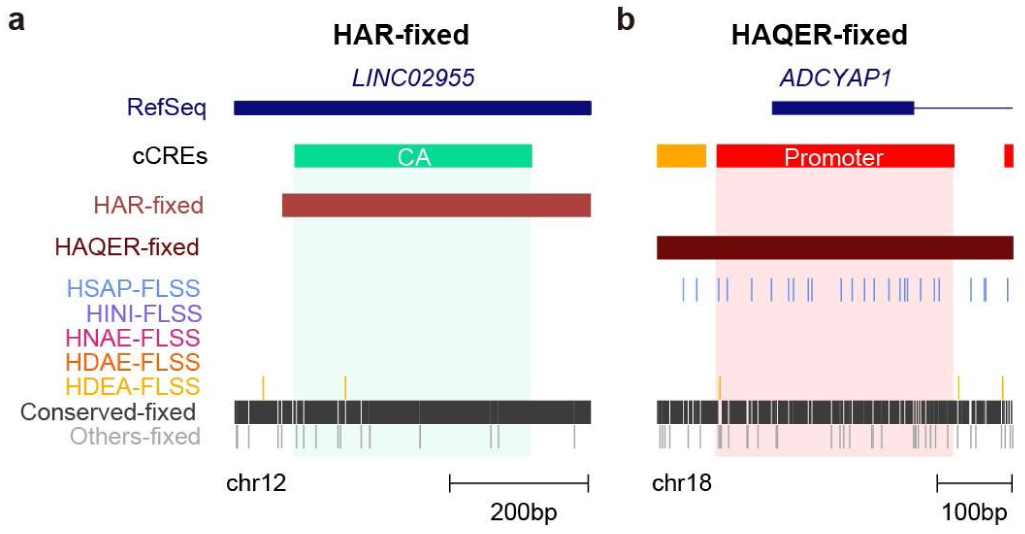
Human-FLSS coverages on the longest CCREs in HAR and HAQER. **a**, Genome browser view of the longest HAR-fixed region overlapping a cCRE, in *LINC02955*. Tracks show the RefSeq gene model, cCRE annotation, HAR-fixed and HAQER-fixed regions, fixed lineage-specific sequence sets for five simian lineages, and positions classified as conserved-fixed or others-fixed in the 9-way alignment. This region does not overlap the HAQER-fixed sequence or human-FLSS. **b**, Genome browser view of the longest HAQER-fixed region overlapping a cCRE, at the *ADCYAP1* promoter. This region does not overlap the HAR-fixed sequence but does overlap multiple human-FLSS positions.

**Extended Data Figure 6.**
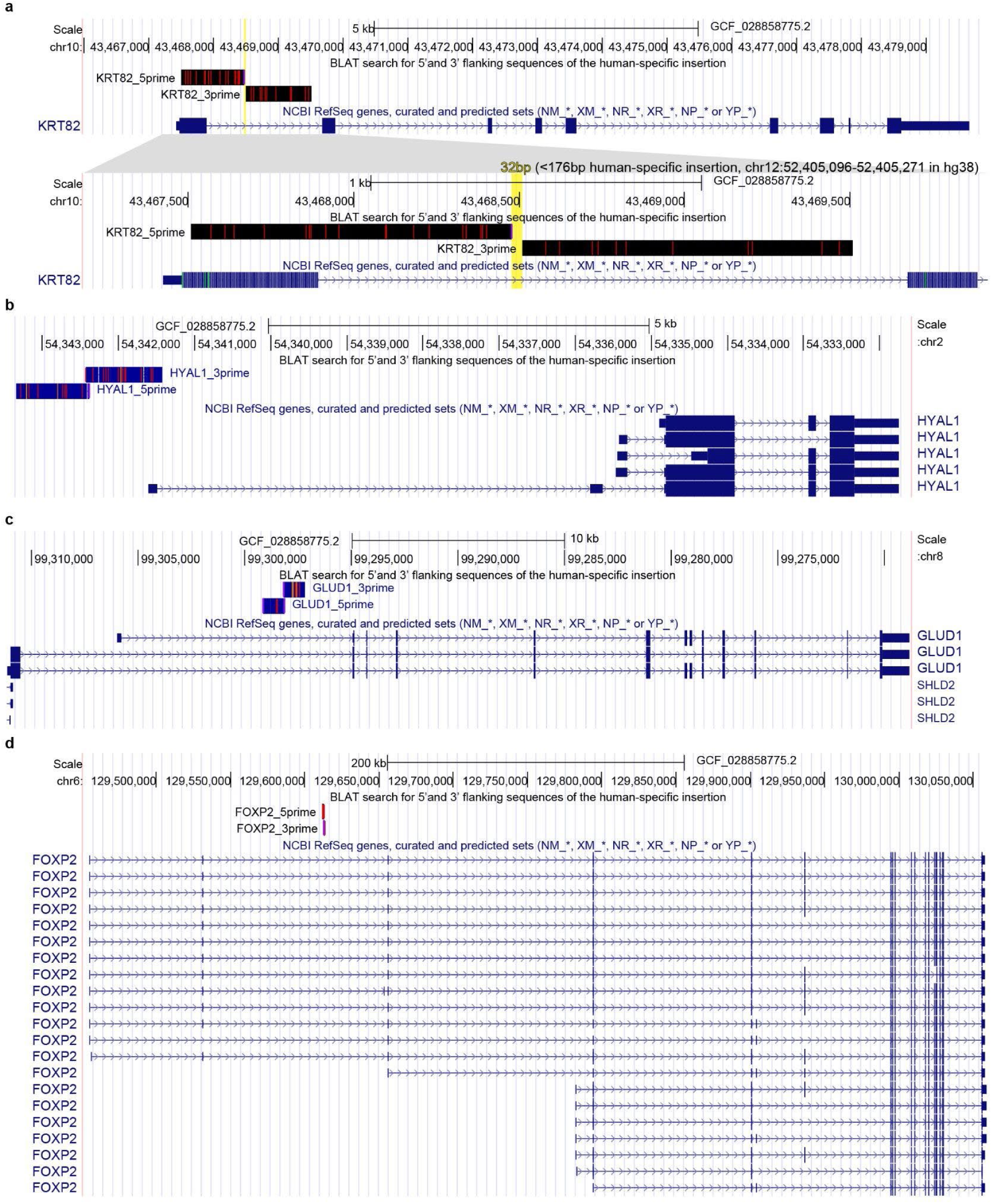
Evidence for absence of selected human-specific insertions in the chimpanzee T2T assembly. **a**, BLAT alignment of sequences flanking the human-specific enhancer EH38E1612606 at *KRT82* against the chimpanzee genome assembly (GCF_028858775.2; chr10). Top, overview of the KRT82 locus at 5-kb scale showing BLAT hits for the 5’ and 3’ flanking sequences together with the NCBI RefSeq gene model. Bottom, zoomed view at 1-kb scale. **b**, BLAT alignment of sequences flanking a human-specific SVA_D insertion at *HYAL1* against the chimpanzee genome assembly (GCF_028858775.2; chr2). The 5’ and 3’ flanking sequences map upstream of *HYAL1* with overlapping hits and no intervening chimpanzee sequence corresponding to the human insertion. **c**, BLAT alignment of sequences flanking the SVA_D-derived enhancer cCRE EH38E2914471 at *GLUD1* against the chimpanzee genome assembly (GCF_028858775.2; chr8). The flanking sequences map upstream of *GLUD1* with overlapping hits and no intervening chimpanzee sequence corresponding to the human insertion. **d**, BLAT alignment of sequences flanking a human-specific SINE insertion overlapping the *FOXP2* enhancer cCRE against the chimpanzee genome assembly (GCF_028858775.2; chr6). The 5’ and 3’ flanking sequences align to a single locus with no intervening chimpanzee sequence corresponding to the human insertion.

**Extended Data Figure 7.**
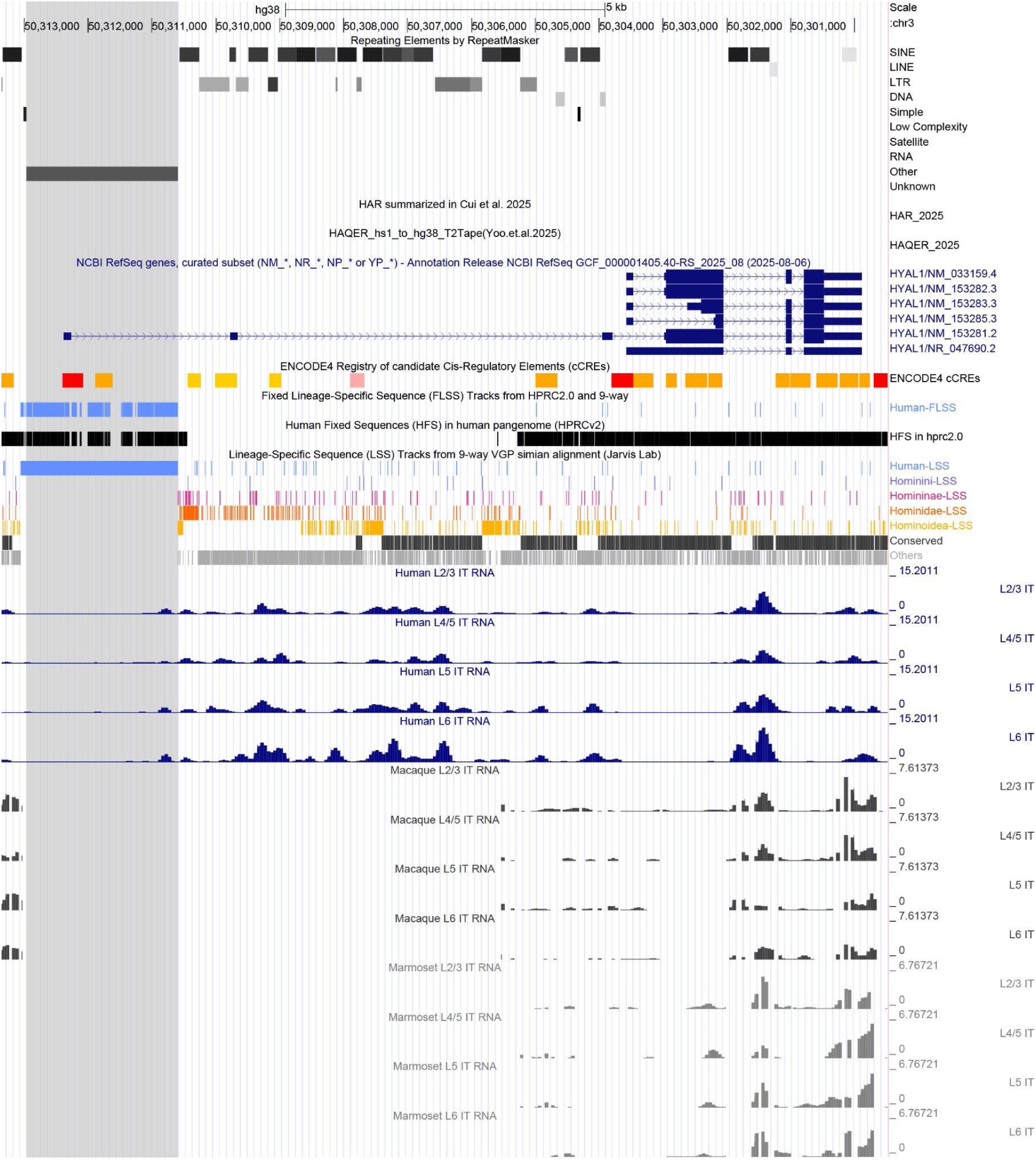
*HYAL1* locus and cross-species IT-neuron expression. UCSC Genome Browser view of the *HYAL1* locus (chr3:50,300,000–50,314,000; hg38), showing RepeatMasker annotation, HAR and HAQER tracks, RefSeq gene models, ENCODE candidate cis-regulatory elements (cCREs), fixed lineage-specific sequence tracks, and single-nucleus RNA-seq expression tracks from intratelencephalic (IT) neuron subtypes (L2/3, L4/5, L5, and L6) in human, macaque, and marmoset. The grey shaded region marks a human-specific SVA_D insertion (https://genome.ucsc.edu/s/clee03/HGSE2026_HYAL1_exp).

**Extended Data Figure 8.**
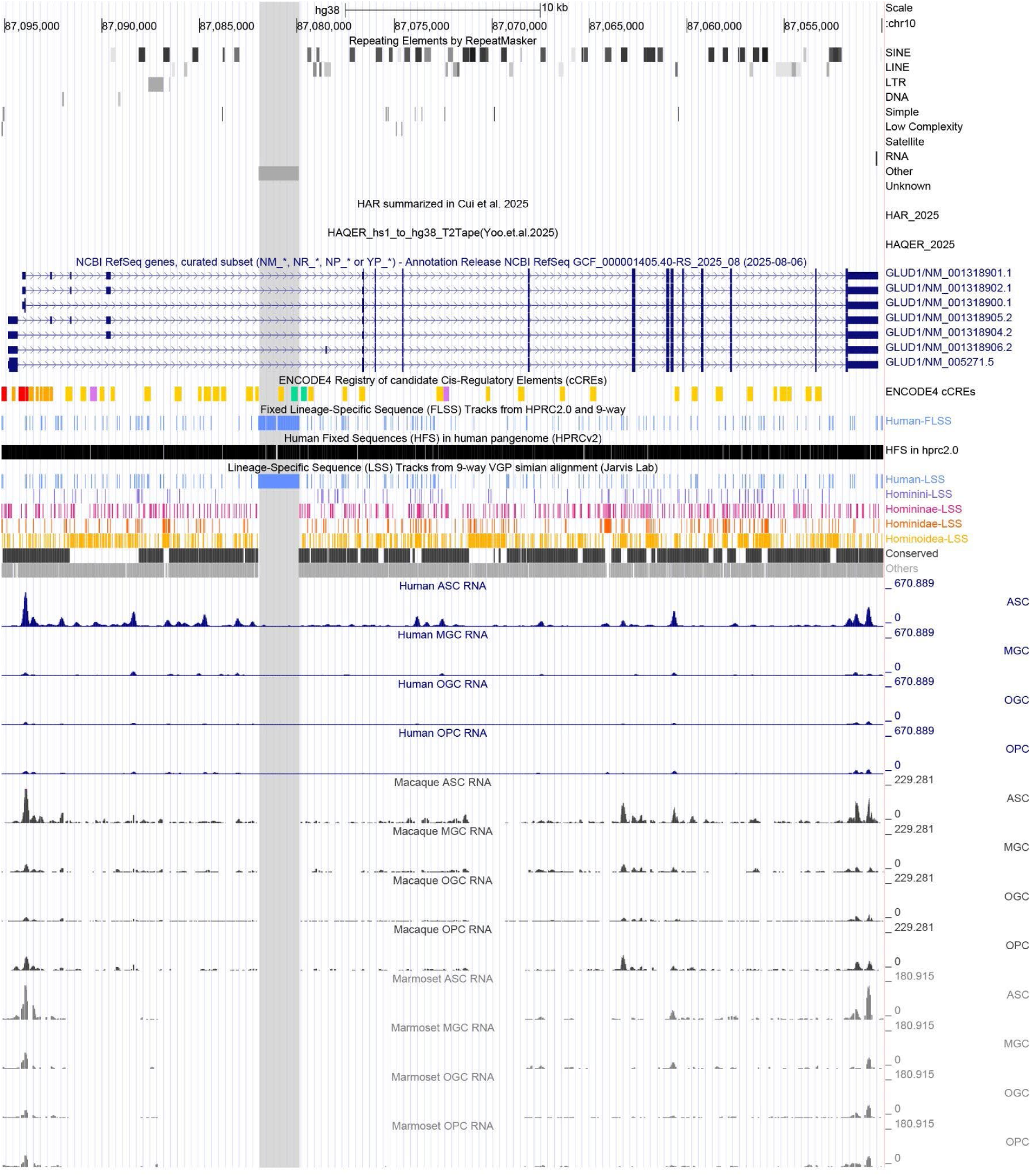
*GLUD1* locus and cross-species expression in glial cell types. UCSC Genome Browser view of the *GLUD1* locus (chr10:87,050,000–87,097,000; hg38), showing RepeatMasker annotation, HAR and HAQER tracks, RefSeq gene models, ENCODE candidate cis-regulatory elements (cCREs), fixed lineage-specific sequence tracks, and single-nucleus RNA-seq expression tracks from glial cell types (astrocytes (ASC), microglia (MGC), oligodendrocytes (OGC), and oligodendrocyte precursor cells (OPC)), in human, macaque, and marmoset (https://genome.ucsc.edu/s/clee03/HGSE2026_GLUD1_exp).

**Extended Data Figure 9.**
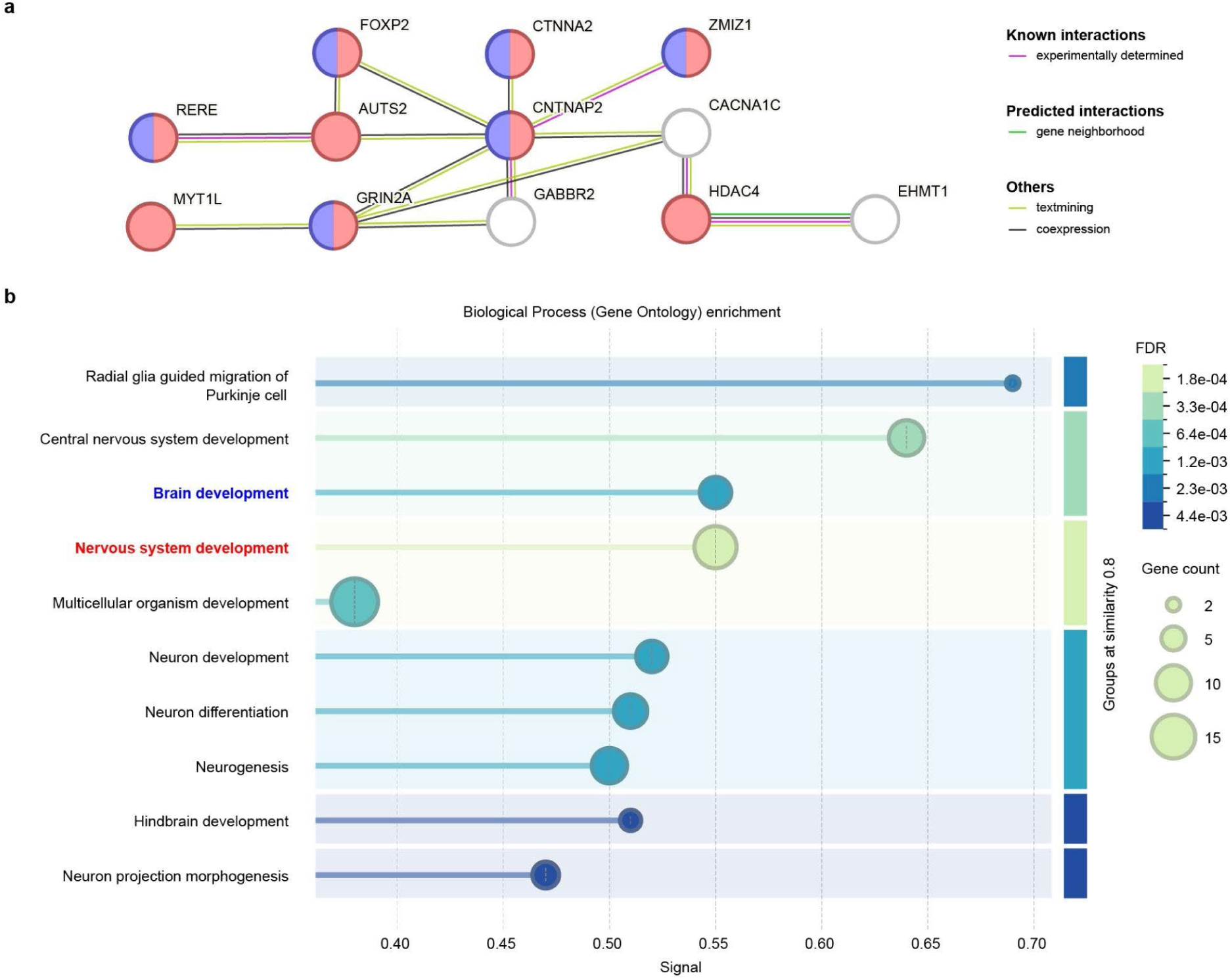
Protein–protein interaction network and functional enrichment of 26 convergent genes. **a**, STRING protein–protein interaction network for 26 genes identified by convergence across human-FLSS ranking, differential expression in human brain, autistic brain differential expression, and the Human Phenotype Ontology terms autistic behavior and language impairment. **b**, Gene Ontology Biological Process enrichment analysis of the same 26-gene set. Terms are grouped by semantic similarity (threshold = 0.8), and the x-axis indicates enrichment signal.

**Extended Data Figure 10.**
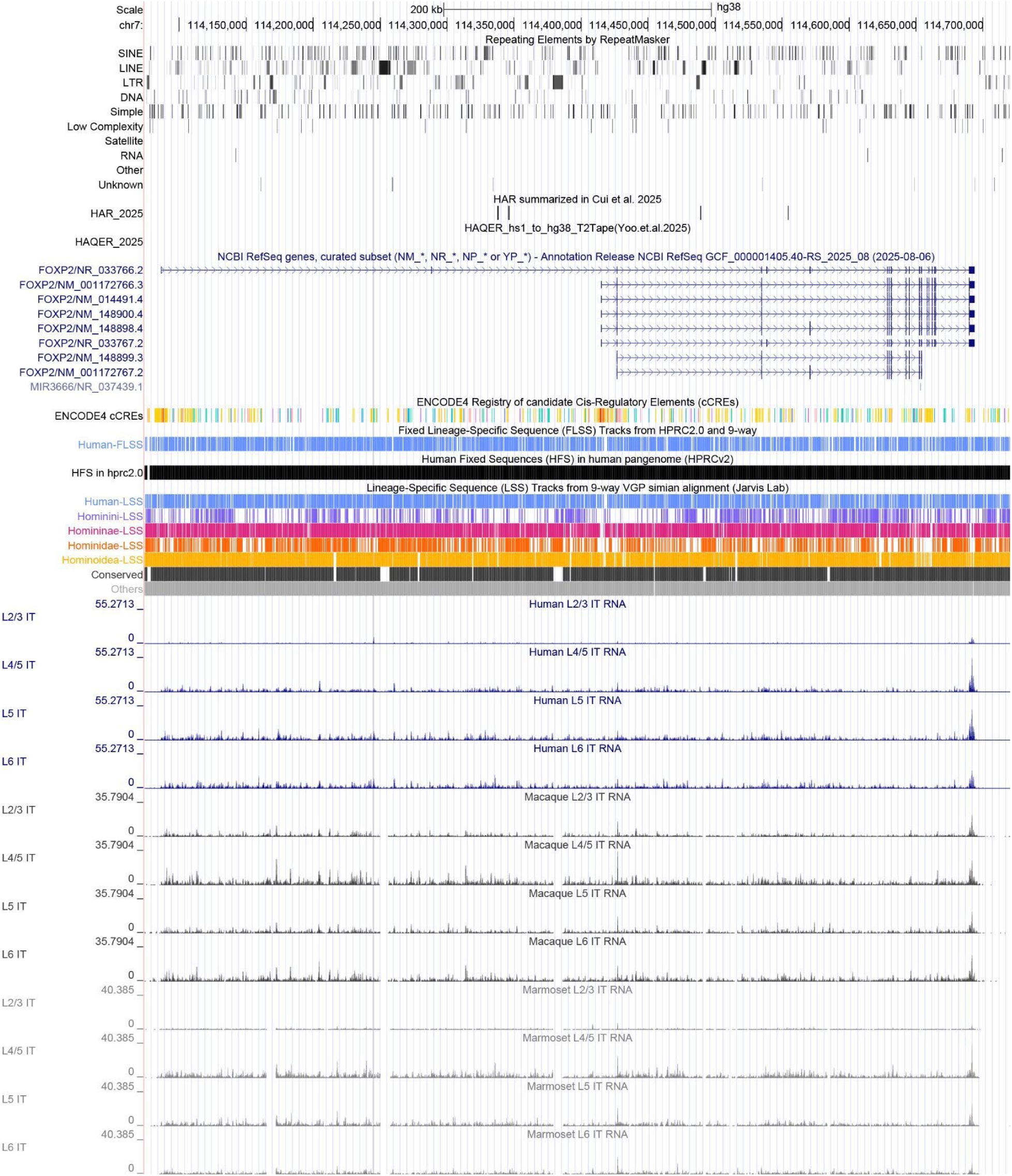
*FOXP2* locus and cross-species IT-neuron expression. UCSC Genome Browser view of the *FOXP2* locus (chr7:114,086,327–114,693,772; hg38), showing RepeatMasker annotation, HAR and HAQER tracks, RefSeq gene models, ENCODE candidate cis-regulatory elements (cCREs), FLSS, and snRNAseq expression tracks from intratelencephalic (IT) neuron subtypes (L2/3, L4/5, L5, and L6) in human, macaque, and marmoset (https://genome.ucsc.edu/s/clee03/HGSE2026_FOXP2_exp).

## Data availability

The RefSeq genomes used in this study are available in NCBI under the accession numbers listed in **Supplementary Tables** (<u>link</u>). The public track hub for Human Genome Sequence Evolution (HGSE, https://hgse2026.s3.us-east-2.amazonaws.com/hub.txt) can be easily accessed for all FLSS catalogs at the UCSC Genome Browser session https://genome.ucsc.edu/s/clee03/HGSE.

## Code availability

The GOtools software is available on GitHub (https://github.com/csgDarwin/gotools). All scripts can be found at (https://github.com/csgDarwin/gotools/tree/main/manuscript/scripts).

## Acknowledgement

We would like to thank the Human Pangenome Reference Consortium (BioProject ID: PRJNA730823), its funder, the National Human Genome Research Institute (NHGRI), and the Vertebrate Genomes Project (BioProject ID: PRJNA489243). We would like to thank the ENCODE project, the RefSeq eukaryote annotation team, the PsychENCODE project, and the UCSC Genome Browser. We would like to thank Dr. Constance Scharff and Dr. Marwa Mahmoud for their insightful comments on this manuscript. This work was financially supported by the NIH T-R01 (R01DC018691 for CL, EDJ) and the Howard Hughes Medical Institute (MHD, EDJ), USA.

## Author information

These authors contributed equally: CSG and MHD

## Contributions

CSG, MHD, and CL conceived the fixed-lineage specific sequence framework. CSG and CL developed GOtools. CL generated the guide tree and the 9-way simian alignment and optimized the 464-way human pangenome alignment as inputs. CSG performed variant and fixed position calling to produce all sequence catalogs. CL generated the HGSE track hub with FLSS catalogs in the UCSC genome browser. CSG and CL validated the variant calling performance of GOtools. CL calculated and visualized proportions of variants and fixation rates in regulatory regions and genes. CL and MD investigated previously reported genes, *FOXP2*, *NOVA1*, and *TBXT*. CL identified hair-related candidate variation in *KRT82*. CL and CSG compared HARs and HAQERs to the human-FLSS catalog. MHD designed and performed comparative snRNA analyses of ape midtemporal gyrus. MHD and CL integrated DEGs and regulatory FLSS. CL confirmed expression patterns using simian primary motor cortex snRNA profiles. MHD and CL described mutations linked to human brain-specialized expression of *HYAL1* and *GLUD1*. CSG, MHD, and CL contributed to gene ontology analyses. CL described the brain expression patterns of autism patients and the protein-protein network of key candidate genes. CL validated human-specific insertions creating regulatory elements in *FOXP2*, *KRT82*, *HYAL1*, and *GLAD1* by scanning the chimpanzee genome. CSG, MHD, and CL drafted the manuscript. All authors reviewed the draft. EDJ secured funding. CL and EDJ co-supervised the study.

## References

1. The Human Condition—A Molecular Approach. Cell 157, 216–226 (2014).

2. Pollen, A. A., Kilik, U., Lowe, C. B. & Camp, J. G. Human-specific genetics: new tools to explore the molecular and cellular basis of human evolution. Nature Reviews Genetics 24, 687–711 (2023).

3. Tubbs, R. S. et al. Enigmatic human tails: A review of their history, embryology, classification, and clinical manifestations. Clin Anat 29, 430–438 (2016).

4. Xia, B. et al. On the genetic basis of tail-loss evolution in humans and apes. Nature 626, 1042–1048 (2024).

5. Pollard, K. S. et al. An RNA gene expressed during cortical development evolved rapidly in humans. Nature 443, 167–172 (2006).

6. Pollard, K. S. et al. Forces Shaping the Fastest Evolving Regions in the Human Genome. PLOS Genetics 2, e168 (2006).

7. Lindblad-Toh, K. et al. A high-resolution map of human evolutionary constraint using 29 mammals. Nature 478, 476–482 (2011).

8. Gittelman, R. M. et al. Comprehensive identification and analysis of human accelerated regulatory DNA. Genome Res 25, 1245–1255 (2015).

9. Girskis, K. M. et al. Rewiring of human neurodevelopmental gene regulatory programs by human accelerated regions. Neuron 109, 3239–3251.e7 (2021).

10. Capra, J. A., Erwin, G. D., McKinsey, G., Rubenstein, J. L. R. & Pollard, K. S. Many human accelerated regions are developmental enhancers. Philos Trans R Soc Lond B Biol Sci 368, 20130025 (2013).

11. Mangan, R. J. et al. Adaptive sequence divergence forged new neurodevelopmental enhancers in humans. Cell 185, 4587–4603.e23 (2022).

12. Yoo, D. et al. Complete sequencing of ape genomes. Nature 641, 401–418 (2025).

13. Luo, Y., et al. Intraspecific sequence variation and complete genomes refine the identification of rapidly evolved regions in humans. bioRxiv (2025) doi:10.1101/2025.10.20.683446.

14. Kim, J. et al. False gene and chromosome losses in genome assemblies caused by GC content variation and repeats. Genome Biol. 23, 204 (2022).

15. Ko, B. J. et al. Widespread false gene gains caused by duplication errors in genome assemblies. Genome Biology 23, 205 (2022).

16. Korlach, J. et al. De novo PacBio long-read and phased avian genome assemblies correct and add to reference genes generated with intermediate and short reads. Gigascience 6, 1–16 (2017).

17. Enard, W. et al. Molecular evolution of FOXP2, a gene involved in speech and language. Nature 418, 869–872 (2002).

18. Tajima, Y. et al. A humanized NOVA1 splicing factor alters mouse vocal communications. Nature Communications 16, 1542 (2025).

19. Moore, J. E. et al. An expanded registry of candidate cis-regulatory elements. Nature (2026) doi:10.1038/s41586-025-09909-9.

20. ENCODE Project Consortium. An integrated encyclopedia of DNA elements in the human genome. Nature 489, 57–74 (2012).

21. Hitz, B. C., et al. The ENCODE Uniform Analysis Pipelines. bioRxiv (2023) doi:10.1101/2023.04.04.535623.

22. Kagda, M. S. et al. Data navigation on the ENCODE portal. Nat Commun 16, 9592 (2025).

23. Erjavec, S. O. et al. Whole exome sequencing in Alopecia Areata identifies rare variants in KRT82. Nat. Commun. 13, 800 (2022).

24. Zhao, B. et al. Single nucleotide polymorphisms in the KRT82 promoter region modulate irregular thickening and patchiness in the dorsal skin of New Zealand rabbits. BMC Genomics 25, 458 (2024).

25. Jorstad, N. L. et al. Comparative transcriptomics reveals human-specific cortical features. Science 382, eade9516 (2023).

26. King, M. C. & Wilson, A. C. Evolution at two levels in humans and chimpanzees. Science 188, 107–116 (1975).

27. Wray, G. A. The evolutionary significance of cis-regulatory mutations. Nat Rev Genet 8, 206–216 (2007).

28. Franchini, L. F. & Pollard, K. S. Human evolution: the non-coding revolution. BMC Biol 15, 89 (2017).

29. Kuhlwilm, M. & Boeckx, C. A catalog of single nucleotide changes distinguishing modern humans from archaic hominins. Sci Rep 9, 8463 (2019).

30. Weiss, C. V. et al. The -regulatory effects of modern human-specific variants. Elife 10, (2021).

31. Uebbing, S. et al. Massively parallel discovery of human-specific substitutions that alter enhancer activity. Proc Natl Acad Sci U S A 118, (2021).

32. Cui, X. et al. Comparative characterization of human accelerated regions in neurons. Nature 640, 991–999 (2025).

33. Lee, Y.-G., Lee, J.-Y., Kim, J. & Kim, Y.-J. Insertion variants missing in the human reference genome are widespread among human populations. BMC Biol 18, 167 (2020).

34. Rhie, A. et al. Towards complete and error-free genome assemblies of all vertebrate species. Nature 592, 737–746 (2021).

35. Larivière, D. et al. Scalable, accessible and reproducible reference genome assembly and evaluation in Galaxy. Nature Biotechnology 42, 367–370 (2024).

36. Nurk, S. et al. The complete sequence of a human genome. Science 376, 44–53 (2022).

37. Liao, W.-W. et al. A draft human pangenome reference. Nature 617, 312–324 (2023).

38. Bakken, T. E. et al. Comparative cellular analysis of motor cortex in human, marmoset and mouse. Nature 598, 111–119 (2021).

39. A multimodal cell census and atlas of the mammalian primary motor cortex. Nature 598, 86–102 (2021).

40. Goldfarb, T. et al. NCBI RefSeq: reference sequence standards through 25 years of curation and annotation. Nucleic Acids Res. 53, D243–D257 (2025).

41. Jorstad, N. L. et al. Transcriptomic cytoarchitecture reveals principles of human neocortex organization. Science 382, eadf6812 (2023).

42. Banerjee-Basu, S. & Packer, A. SFARI Gene: an evolving database for the autism research community. Dis Model Mech 3, 133–135 (2010).

43. Wamsley, B. et al. Molecular cascades and cell type-specific signatures in ASD revealed by single-cell genomics. Science 384, eadh2602 (2024).

44. Szklarczyk, D. et al. The STRING database in 2023: protein-protein association networks and functional enrichment analyses for any sequenced genome of interest. Nucleic Acids Res 51, D638–D646 (2023).

45. McLean, C. Y. et al. Human-specific loss of regulatory DNA and the evolution of human-specific traits. Nature 471, 216–219 (2011).

46. Xue, J. R. et al. The functional and evolutionary impacts of human-specific deletions in conserved elements. Science 380, eabn2253 (2023).

47. Lee, C., Davenport, M. H. & Jarvis, E. D. A human specific CCG repeat in the RBFOX1 promoter is implicated in speech and autism. bioRxiv (2026) doi:10.64898/2026.04.21.719679.

