## Supplementary material for "Fixed human pangenome sequences reveal origins of common human traits": Online Methods

#### Genome-based guide tree inference for cross-species alignment

A genome-based guide tree for the 9-way simian whole-genome alignment was inferred using ROADIES[1] (v0.1.10), installed with Conda (conda install roadies=0.1.10). The analysis included the human reference genome GRCh38.p14/hg38 (GCF_000001405.40) and eight simian reference assemblies[2,3]: chimpanzee mPanTro3 (GCF_028858775.2), bonobo mPanPan1 (GCF_029289425.2), gorilla mGorGor1 (GCF_029281585.2), Sumatran orangutan mPonAbe1 (GCF_028885655.2), Bornean orangutan mPonPyg2 (GCF_028885625.2), siamang mSymSyn1 (GCF_028878055.3), crab-eating macaque MFA8 (GCF_037993035.2), and common marmoset calJac240 (GCA_049354715.1). Each assembly was provided as a single FASTA file with chromosomal scaffolds.

No reference tree was provided (REFERENCE=NULL). The analysis used 500 bp sampled genomic segments (LENGTH=500), 250 sampled segments per iteration (GENE_COUNT=250), a minimum uppercase sequence fraction of 0.90 (UPPER_CASE=0.90), LASTZ alignment thresholds of 65% identity, 85% coverage, and 85% continuity (IDENTITY=65, COVERAGE=85, CONTINUITY=85), a maximum of 10 duplicated homologous copies per genome (MAX_DUP=10), one LASTZ sampling step (STEPS=1), PASTA filtering parameters of FILTERFRAGMENTS=0.50 and MASKSITES=0.02, a convergence support threshold of 0.95 (SUPPORT_THRESHOLD=0.95), and one parallel ROADIES instance (NUM_INSTANCES=1). The run used MIN_ALIGN=4, and deep phylogeny mode was not enabled.

Because the input assemblies were multi-gigabase simian genomes, LASTZ was run using lastz_40[4] (v1.04.22) compiled with 63-bit sequence indexing for 2-4 Gb genome sizes. Multiple sequence alignments were filtered, and gene trees were inferred with RAxML-NG[5] under a GTR+G+F model. The resulting gene trees were summarized with ASTRAL-Pro3[6] (v1.19.3.6). The run completed all 772 Snakemake jobs and converged after the first iteration, generating 247 gene trees. The final ASTRAL-Pro3 species tree had a score of 27,040 across 30,483 equivalent quartets and was used as the guide phylogeny for downstream 9-way simian whole-genome alignment:

*((((((hg38:0.02070700,(mPanTro3:0.00354400,mPanPan1:0.00230000):0.00281200):0.00168400,mGorGor1:0.00763300):0.00758300,(mPonAbe1:0.00025100,mPonPyg2:0.00058300):0.01676400):0.00248900,mSymSyn1:0.03029800):0.01466500,MFA8:0.03369700):0.04939000,calJac240:0.04939000);*

**9-way cross-species simian genome-wide alignment**

Using the ROADIES-inferred genome-based species tree as the guide phylogeny, we generated a 9-way simian whole-genome alignment with Progressive Cactus[7] (v3.0.1). The input assemblies were the same nine assemblies used for guide-tree inference. Each assembly was provided as a single FASTA file, and the ROADIES-derived rooted species tree was specified in the Cactus sequence file, with the common marmoset as the most distant outgroup.

Progressive Cactus was executed through Toil using 30 CPU cores, generating the hierarchical alignment file '9way_primates.hal'. For downstream hg38-referenced analyses, the completed HAL alignment was exported to a single MAF file using ***cactus-hal2maf***[7]. The conversion used hg38 as the reference genome, --filterGapCausingDupes, --outType single, 500-kb reference chunks, 30 batch cores, ancestral sequences excluded, caching disabled, a dedicated working directory, and a maximum disk allocation of 500 GB. This generated '9way_primates_hg38.single.maf.gz'.

For UCSC Genome Browser visualization, the single MAF file was converted to bigMaf format using ***cactus-maf2bigmaf***[7], with '9way_primates.hal' supplied as the source HAL file. The workflow obtained hg38 chromosome sizes using halStats, filtered duplicate and low-quality MAF records using *mafDuplicateFilter* and *mafFilter*, converted the filtered alignments using ***mafToBigMaf***[8], and indexed the output using ***bedToBigBed***[8]. A summary bigBed file was generated with ***hgLoadMafSummary***[8]. The final browser-ready outputs were '9way_primates_hg38.single.bigmaf.bb' and '9way_primates_hg38.single.bigmaf.summary.bb'.

**464-way human-pangenome**

The within-species human pangenome alignment was generated in collaboration with the Human Pangenome Reference Consortium v2.0 using Minigraph-Cactus[9]. The precomputed HAL file, 'hprc-v2.0-mc-chm13.full.hal (link)', was downloaded from the HPRC Data Explorer (42basepairs) and used as the source alignment for assessing population-level sequence invariance across human haplotypes. For downstream hg38-referenced analyses, the HAL alignment was exported to a singleton MAF file using ***cactus-hal2maf***[7] with hg38 as the reference genome, --outType single, --filterGapCausingDupes, 100-kb reference chunks, 30 batch cores, and ancestral sequences excluded. This generated 'hprc-v2.0-mc-hg38.singleton_rename.maf.gz'. For UCSC Genome Browser visualization, the singleton MAF file was converted to bigMaf format using ***cactus-maf2bigmaf***[7], with the original HPRCv.2.0 HAL file supplied for reference genome information and --maxDisk 1000G, producing 'hprc-v2.0-mc-hg38.singleton_rename.bigmaf.bb'. The resulting hg38-referenced pangenome alignment was used for a new pangenome-aware framework by integrating the 9-way simian alignment to identify lineage-specific sequences fixed across the sampled human pangenome in the next step.

**FLSS identification using GOtools**

We used ***go2var*** and ***go2fix*** from the GOtools Python package (https://github.com/csgDarwin/GOtools) to investigate human genome sequence evolution in both the 9-way simian alignment and the 464-way human pangenome alignment.

First, we ran ***go2var*** on the 9-way simian alignment ('9way_primates_hg38.single.maf.gz') to identify lineage-specific sequences (LSS) for each monophyletic lineage including human: *Homo sapiens* (humans: hg38), Hominini (humans and *pan* lineage: hg38, mPanTro3, mPanPan1),

Homininae (African great apes: hg38, mPanTro3, mPanPan1, mGorGor1), Hominidae (great apes: hg38, mPanTro3, mPanPan1, mGorGor1, mPonAbe1, mPonPyg2), and Hominoidea (apes: hg38, mPanTro3, mPanPan1, mGorGor1, mPonAbe1, mPonPyg2, mSymSyn1). For each lineage, ***go2var*** was run with a lineage-specific JSON configuration file specifying the required target species and alignment order. A nucleotide site was called a variant for a given lineage when the target species shared the same base, and every other species carried a different base. We repeated this test across five lineage categories, retaining only homologous sites with an assembly-gap-free hg38 base (no 'N'). Each variant site was output to its corresponding '[lineage name]-LSS.bed' file (e.g., 'Human-LSS.bed').

Additionally, we identified fully conserved sites across the 9-way alignment using ***go2var***. Fully conserved sites were defined as homologous hg38 sites at which all aligned species carried the same A, C, G, or T base. To define the other remaining sites in the human reference genome, we merged all lineage-specific BED files ([lineage name]-LSS.bed) with the fully conserved BED file (conserved.bed) using ***merge*** in bedtools[10] (v2.31.1). We then subtracted these merged intervals from the hg38 genome, restricted to positions with A, C, G, or T bases, using ***bedtools subtract***[10]. The resulting intervals were classified as "others" sites, representing hg38 positions that were neither lineage-specific nor fully conserved in the 9-way alignment.

Next, we ran ***go2fix*** on the HPRC pangenome alignment ('hprc-v2.0-mc-hg38.singleton_rename.maf.gz') to identify positions fixed across human haplotypes. Because haplotype coverage varies among autosomes and sex chromosomes, we first counted the maximum number of haplotypes covered in HPRCv2 alignment blocks for each class (464 for autosomes, 363 for chromosome X, and 118 for chromosome Y). We then ran ***go2fix -m*** separately for each chromosome class, setting the per-class maximum (464 for autosomes, 363 for chromosome X,

118 for chromosome Y) to require fixation across the full set of covered haplotypes within each class. If a homologous site contained the maximum number of haplotypes within its chromosome class and all haplotypes shared the same base, it was considered a fixed site and written to the 'HFS.bed' file. Finally, we defined fixed lineage-specific sites (FLSS) as the intersection of 'LSS.bed' of all five lineages and 'HFS.bed', computed with ***bedtools intersect***[10]. Human-FLSS were cross-checked on autosomes using ***maf2var*** from ReAlignPro[11] for variant-calling performance, and the final FLSS catalogs from all lineages were manually assessed across all chromosomal scaffolds in a new public track hub in the UCSC Genome Browser.

**FLSS catalogs in a public UCSC track hub**

To facilitate manual validation and public visualization of the sequence-evolution catalogs, we built a publicly accessible UCSC Track Hub for Human Genome Sequence Evolution (HGSE, v2026). BED-based intermediate and final datasets, including LSS, HFS, and FLSS catalogs, were converted to bigBed format using ***bedToBigBed***[8]. These annotation tracks, along with the 9-way simian and 464-way human pangenome alignment tracks, were uploaded to an Amazon Web Services (AWS)-hosted track hub (https://hgse2026.s3.us-east-2.amazonaws.com/hub.txt) for visualization in the UCSC Genome Browser[8].

**Regulatory FLSS on human regulatory elements**

To characterize regulatory functions of FLSS in the human genome, we downloaded the human candidate cis-regulatory elements (cCREs; https://downloads.wenglab.org/Registry-V4/GRCh38-cCREs.bed) for 2,341,591 loci from the SCREEN Portal (https://screen.wenglab.org/) in the ENCODE project[12–15]. These cCREs were split into regulatory classes containing pELS, dELS,

PLS, *CTCF*-only, and DNase-H3K4me3, and intersected with the FLSS of each lineage to define regulatory FLSS using ***bedtools intersect***[10]. After that, the proportions of regulatory FLSS per cCRE regions were calculated using ***bedtools coverage***[10]**,** and FLSS were filtered using a cutoff (>25%) to identify lineage-specific and human-fixed regulatory elements (e.g., human-specific and fixed cCREs). To support a key example gene, *KRT82*, its tissue-dependent expression profiles from bulk RNA-seq were searched and visualized in the SCREEN (https://screen.wenglab.org/GRCh38/gene/KRT82?open=BYegjCDSBKAqAcAmEAGIA).

**FLSS on human promoter and gene bodies**

To characterize the genic functions of FLSS, the human RefSeq gene annotation for 59,046 genes (v2.2; Release RS_2025_08, GRCh38.p14, GCF_000001405.40) was retrieved in GTF format from the NCBI RefSeq database[16]. We assigned putative promoter intervals to the 1 kb 5′ upstream window of each annotated gene using ***AddPro*** in GOtools and combined the promoter and gene-body intervals into a single gene annotation set. We used ***bedtools intersect***[10] to intersect regulatory FLSS in cCREs with the promoter-extended gene annotation, identifying regulatory FLSS within each gene. Then we calculated the number of regulatory FLSS per gene and ranked genes by this count to identify the top 1,000 genes across the five key lineages: human, hominini, homininae, hominidae, and hominoidea.

**Proportion matrices of human genome sequence evolution**

To generate genome-wide proportion matrices, an in-house Python pipeline was developed based on ***bedtools intersect***[10] and base-pair coverage summarization. All BED-like input files were first normalized to BED3 format, filtered to remove header lines and invalid intervals, and sorted by chromosome and start coordinate. The input sequence-evolution categories included LSS for the Homo sapiens, Hominini, Homininae, Hominidae, and Hominoidea lineages; 9-way conserved sites; the other sites; and HFS. Fixed sequence-evolution categories were generated by intersecting each sequence-evolution BED file with the HFS BED file. We then computed pairwise intersections between sequence-evolution categories, fixed sequence-evolution categories, HFS, cCRE annotations, and RefSeq gene annotation sets using bedtools intersect.

Proportions were calculated by summing the base-pair lengths of BED intervals. The denominator for genome-wide proportions was the hg38 callable genome space restricted to A, C, G, and T bases, excluding ambiguous assembly-gap bases. The denominator for feature-specific proportions was the total base-pair length of each regulatory or gene annotation category. Regulatory categories included all cCREs, PLS, pELS, dELS, CA-TF, CA-CTCF, CA-H3K4me3, CA, and TF elements. Gene annotation categories included RefSeq gene bodies, 5' upstream 1 kb regions, 3' downstream 1 kb regions, promoters (±1kb from transcription start sites), core promoters (±150 from transcription start sites), introns, exons, coding regions, 5′ UTRs, 3′ UTRs, and intergenic regions.

For each feature category, we calculated the genome-wide fraction occupied by the feature, the fraction of the feature overlapped by each sequence-evolution category, and the fixation rate for each category, defined as the number of fixed base pairs in that category within the feature divided by the total base pairs in that category within the feature. The resulting matrices were exported as TSV files for downstream visualization.

#### Fixation patterns of HAR, HAQER, and Human-LSS

To compare our annotations with previously reported candidate regions related to human evolution, Human Accelerated Regions (HARs)[17–23] and Human Ancestor Quickly Evolved Regions (HAQERs)[24] were retrieved. HAQER coordinates were projected from the hs1 assembly to the hg38 assembly using the UCSC Genome Browser ***LiftOver***[8] tool. Of the original HAQER loci, 300 failed to lift over because the corresponding intervals were fully or partially deleted or split in hg38, and 2,747 HAQER loci were retained in hg38 for downstream analysis. The 3,257 HAR loci and 2,747 lifted HAQER loci in hg38 were intersected with human-LSS, HFS, and human-FLSS annotations using ***bedtools intersect***[10], and the fraction of each HAR or HAQER interval covered by each annotation was calculated using ***bedtools coverage***[10]. To visually compare the features, the entire set of intersection outputs from the HAR and HAQER loci was converted to bigBed format using ***bedToBigBed***[8] and uploaded to the HGSE2026 track hub (https://hgse2026.s3.us-east-2.amazonaws.com/hub.txt).

For visualization and summary statistics, the final column of each ***bedtools coverage***[10] output, representing the fraction of each HAR or HAQER interval covered by the queried annotation, was used. Analyses were restricted to chromosomal scaffolds, chr1–chr22, chrX, and chrY. Non-primary contigs, missing values, non-numeric entries, and values outside the 0–1 range were excluded. Coverage-fraction distributions were visualized using in-house R scripts with ***ggplot2***, after kernel density estimation with the ***density*** function (bw = "nrd0", adjust = 1). For each comparison, we calculated the number and percentage of HAR or HAQER loci below and above the predefined cutoff values, 3.892% for human-LSS-associated comparisons and 0.901% for human-FLSS-associated comparisons. Summary statistics were exported as tab-delimited TSV format.

Next, to shift the focus from genome-scale to regulatory element-scale, we integrated three human-lineage feature sets (HAR, HAQER, and human-FLSS) with the ENCODE cCREs (GRCh38, registry v4). Using ***bedtools intersect -c***[10], we counted, per cCRE, the number of overlapping human-FLSS and the base-pair overlap with HAR-fixed and HAQER-fixed sequences. For each cCRE, we computed a base-pair coverage percentage with respect to each feature set: pct_human_FLSS = human-FLSS_count / ccre_length × 100, pct_HAR_fixed = HAR_fixed_overlap_bp / ccre_length × 100, and pct_HAQER_fixed analogously.

We summarized each annotation class with two complementary metrics: (i) the number of cCREs with at least one base-pair overlap with the feature, expressed as a count and as a percentage of all 2,348,854 cCREs; and (ii) the unweighted mean of the per-cCRE metric (coverage percentage for human-FLSS, HAR-fixed, and HAQER-fixed), computed across all 2,348,854 cCREs, including those with zero overlap.

We performed visualization in **R** using ***ggplot2***. The number of cCREs with at least one variant overlap was plotted as a barplot across three categories (HAR-fixed, HAQER-fixed, Human-FLSS). For the number of cCREs with at least one overlap, we rendered the three categories as a vertical barplot ordered as HAR-fixed, HAQER-fixed, and Human-FLSS. The y-axis was log10-transformed (range $5 \times 10^2$ to $5 \times 10^6$) to accommodate the wide dynamic range across categories.

**Comparative Midtemporal Gyrus snRNAseq**

Count matrices and metadata tables of preannotated single nucleus transcriptomes from human, chimpanzee, and gorilla midtemporal gyrus were downloaded (https://data.nemoarchive.org/publication_release/Great_Ape_MTG_Analysis/), read into R, randomly subset to 50k nuclei per species, converted to human orthologous gene_ids, and merged to create a 150k nuclei, mixed species Seurat object. We separately compared the expression of each human cell type to homologous chimpanzee and gorilla cell types within this object using Seurat::FindMarkers, retaining only those results that were consistently differential, meaning the same direction of effect compared to the homologous cells from either ape. From this list of filtered results tables, we constructed a matrix of human-differential-direction by gene by cell-type. We hierarchically clustered this matrix (hclust) to identify gene sets with consistent differential expression patterns (cutree, k = 1000), then filtered to sets with >100 genes. The table of cluster assignment by gene was then joined with our measurements of % human FLSS to identify genes with human-specific regulatory features. Single-cell transcriptome profiles of the primary motor cortex of human, macaque, and marmoset were searched and visualized in the UCSC track hub, CCG2025[11] (https://ccg2025.s3.us-east-2.amazonaws.com/hub.txt).

**Functional enrichment of top-ranked genes**

We performed functional enrichment analysis with g:Profiler[25] using the gprofiler-official Python client (v1.0.0; database release e113_eg59_p19). We queried the top 1,000 genes per lineage against three reference databases: Gene Ontology Cellular Component[26-27](GO:CC; release 2025-03-16), KEGG[28,29] (release 2024-01-22), and the Human Phenotype Ontology[30] (HP; release 2025-05). We assessed significance with the false discovery rate method at a 0.05 threshold (Benjamini–Hochberg). Before enrichment, we resolved ambiguous gene symbols by selecting the Ensembl gene ID with the most GO annotations, replicating the default options of the g:Profiler web interface. We exported results from all sources as a single CSV per lineage, then combined them into a single table to generate heatmap plots.

Across the five lineages (Human, Hominini, Homininae, Hominidae, Hominoidea), we computed per-term fold enrichment from the g:Profiler output as FE = (observed/expected) gene overlap and log2-transformed the values. For each annotation source separately (GO:CC, KEGG, HP), we ranked terms within each lineage by adjusted *p*-value. For the GO:CC and KEGG analyses, we retained the union of the top N terms per lineage (N = 20) for plotting. For the HP analysis, we retained all terms. We grouped retained terms by the lineages in which they were enriched. Next, we ordered groups with the most broadly shared first, then by lineage position within each group. Within each group, we sorted terms by their minimum adjusted *p*-value across lineages, placing the most significant terms at the top. We coloured heatmap cells by log2 fold enrichment.

**Single-cell expression analysis for ASD patient brains**

To examine single-cell expression patterns in autism spectrum disorder (ASD) brains, we narrowed down key candidate genes by intersecting the following features. First, a gene pull was constructed by taking the union of ASD-risk, language-related, and human brain-specialized genes, including differentially expressed genes between human and chimpanzee or gorilla brains (n = 10,233) and SFARI score 1 or syndromic ASD genes (n = 435), along with GO-HP genes annotated with 'autistic behavior' (n = 63), and GO-HP genes annotated with 'language impairment' (n = 90) enriched in the top 1,000 genes with regulatory human-FLSS. This union contained 10,608 genes. Next, we intersected this gene set with the associated, yielding 26 genes for single-cell expression visualization.

For each of the 26 genes, cell-subtype-level differential expression values were extracted from the ASD versus control comparison. The input table from the ASD-snGENE portal[31] (http://solo.bmap.ucla.edu/asdscgene/) contained gene symbols, annotated cell subtypes, log2 fold change values for ASD versus control brains, and FDR-adjusted *p*-values. Cell subtypes were grouped into major classes, including astrocytes, blood-brain barrier-associated cells, excitatory neurons, inhibitory neurons, microglia, oligodendrocytes, oligodendrocyte precursor cells, and other cell types. Differential expression patterns were visualized as a bubble plot using an in-house R script with ***tidyverse***[32], ***ggplot2***[33], and ***scales***[34]. Bubble color represents log2 fold change in ASD versus control brains, with positive values indicating increased expression in ASD and negative values indicating decreased expression. The color scale was centered at zero and truncated between −0.5 and 0.5. Bubble size represents −log10(FDR), capped at 10 to limit the influence of extremely small adjusted *p* values. Differential expression signals that passed an FDR threshold of 0.05 were visually highlighted with an additional outline, although the analysis was primarily used for exploratory visualization rather than to define a new disease-gene set.

**STRING database analysis**

To examine whether the 26 candidate genes formed a functionally connected gene network, we performed protein-protein interaction and functional enrichment analyses using the STRING database[35] (https://string-db.org/). The above 26 key candidate genes were submitted to STRING with Homo sapiens selected as the reference organism. Protein-protein interaction networks were generated using STRING evidence channels, including experimentally determined, predicted, text-mining, and co-expression-based interactions. Network edges were visualized by supporting evidence type. Functional enrichment analysis for these 26 genes was performed within STRING using Gene Ontology Biological Process annotations[35]. Enriched terms were ranked by STRING enrichment signal and filtered using FDR-adjusted significance values. For each enriched term, gene count and FDR-adjusted significance were displayed as bubble size and color, respectively.

### Online Method References

1. Gupta, A., Mirarab, S. & Turakhia, Y. Accurate, scalable, and fully automated inference of species trees from raw genome assemblies using ROADIES. *Proc. Natl. Acad. Sci. U. S. A.* **122**, e2500553122 (2025).
2. Yoo, D. *et al.* Complete sequencing of ape genomes. *Nature* **641**, 401–418 (2025).
3. Giulio Formenti*, Dominic Absolon, Linelle Abueg, Francine Ackerman, Agostinho Antunes, Jean-Nicolas Audet, Jennifer Balacco, Alan Beavan, Mark Blaxter, Thomas Brown, George Butler, Shuo Cao, Eduardo Charvel, Ying Chen, Claudio Ciofi, Hiram Clawson, Helena Conceição, Melanie Couture, Andrew Crawford, Hugues Roest Crollius, Minoli Daigavane, H William Detrich III, Richard Durbin, Erick Duarte, Olivier Fedrigo, Patrick Flannery, Pedro Galante, Erik Garrison, M. P. Thomas Gilbert, François Giudicelli, Simone M. Gable, Andrea Guarracino, Anshu Gupta, Bettina Haase, Michael Hiller, Paul Hime, Kathleen Horan, Kerstin Howe, Danny Jackson, Nivesh Jain, Vinita Joardar, Lin Kang, Byung June Ko, Bonhwang Koo, Sergey Koren, Delphine Lariviere, Chul Lee, Young-Ho Lee, Harris Lewin, Daven Lim, Nicolas Lou, Kateryna Makova, Kirsty McCaffrey, Shane McCarthy, Laia Marín-Gual, Jack A. Medico, Pawel Michalak, Siavash Mirarab, Phillip A. Morin, Jacquelyn Mountcastle, Terence Murphy, Eugene W. Myers, Anton Nekrutenko, Mary J. O’Connell, Brian O'Toole, Benedict Paten, Sadye Paez, Michael Paulini, Sarah Pelan, Andreas Pfenning, Adam M. Phillippy, Martin Pippel, Brendan J. Pinto, Arang Rhie, Aurora Ruiz-Herrera, Yana Safonova, Camilla A. Santos, Michael C. Schatz, Simona Secomandi, Lauren Shalmiyev, Jeramiah Smith, Marco Sollitto, Cibele G. Sotero-Caio, Peter Sudmant, Samuel Talbot, Françoise Thibaud-Nissen, Emma Teeling, Tatiana Tilley, Nataliya Timoshevskaya, Patrick Traore, Yatish Turakhia, Marcela Uliano-Silva, Sonja C. Vernes,

Conor Whelan, Sylke Winkler, Melissa A. Wilson, Jonathan M. D. Wood, Guoyan Zhao, Erich D. Jarvis*, and the Vertebrate Genomes Project Consortium Phase#. The Vertebrate Genomes Project Phase I: A global reference genome resource. *preparing for submission*.

4. Harris, R. S. *Improved Pairwise Alignment of Genomic DNA*. (2007).
5. Kozlov, A. M., Darriba, D., Flouri, T., Morel, B. & Stamatakis, A. RAxML-NG: a fast, scalable and user-friendly tool for maximum likelihood phylogenetic inference. *Bioinformatics* **35**, 4453–4455 (2019).
6. Zhang, C., Nielsen, R. & Mirarab, S. ASTER: A Package for Large-Scale Phylogenomic Reconstructions. *Mol Biol Evol* **42**, (2025).
7. Armstrong, J. *et al.* Progressive Cactus is a multiple-genome aligner for the thousand-genome era. *Nature* **587**, 246–251 (2020).
8. Casper, J. *et al.* The UCSC Genome Browser database: 2026 update. *Nucleic Acids Res* **54**, D1331–D1335 (2026).
9. Hickey, G. *et al.* Pangenome graph construction from genome alignments with Minigraph-Cactus. *Nat. Biotechnol.* **42**, 663–673 (2024).
10. Quinlan, A. R. & Hall, I. M. BEDTools: a flexible suite of utilities for comparing genomic features. *Bioinformatics* **26**, 841–842 (2010).
11. Lee, C., Davenport, M. H. & Jarvis, E. D. A human specific CCG repeat in the RBFOX1 promoter is implicated in speech and autism. *bioRxiv* (2026) doi:10.64898/2026.04.21.719679.
12. ENCODE Project Consortium. An integrated encyclopedia of DNA elements in the human genome. *Nature* **489**, 57–74 (2012).
13. Hitz, B. C. *et al.* The ENCODE Uniform Analysis Pipelines. *bioRxiv* (2023)

doi:10.1101/2023.04.04.535623.

14. Kagda, M. S. *et al.* Data navigation on the ENCODE portal. *Nat Commun* **16**, 9592 (2025).

15. Moore, J. E. *et al.* An expanded registry of candidate cis-regulatory elements. *Nature* (2026) doi:10.1038/s41586-025-09909-9.

16. Goldfarb, T. *et al.* NCBI RefSeq: reference sequence standards through 25 years of curation and annotation. *Nucleic Acids Res.* **53**, D243–D257 (2025).

17. Pollard, K. S. *et al.* An RNA gene expressed during cortical development evolved rapidly in humans. *Nature* **443**, 167–172 (2006).

18. Pollard, K. S. *et al.* Forces Shaping the Fastest Evolving Regions in the Human Genome. *PLOS Genetics* **2**, e168 (2006).

19. Lindblad-Toh, K. *et al.* A high-resolution map of human evolutionary constraint using 29 mammals. *Nature* **478**, 476–482 (2011).

20. Gittelman, R. M. *et al.* Comprehensive identification and analysis of human accelerated regulatory DNA. *Genome Res* **25**, 1245–1255 (2015).

21. Girskis, K. M. *et al.* Rewiring of human neurodevelopmental gene regulatory programs by human accelerated regions. *Neuron* **109**, 3239–3251.e7 (2021).

22. Cui, X. *et al.* Comparative characterization of human accelerated regions in neurons. *Nature* **640**, 991–999 (2025).

23. Capra, J. A., Erwin, G. D., McKinsey, G., Rubenstein, J. L. R. & Pollard, K. S. Many human accelerated regions are developmental enhancers. *Philos Trans R Soc Lond B Biol Sci* **368**, 20130025 (2013).

24. Mangan, R. J. *et al.* Adaptive sequence divergence forged new neurodevelopmental enhancers in humans. *Cell* **185**, 4587–4603.e23 (2022).

25. Kolberg, L. *et al.* g:Profiler-interoperable web service for functional enrichment analysis and gene identifier mapping (2023 update). *Nucleic Acids Res.* **51**, W207–W212 (2023).

26. Ashburner, M. *et al.* Gene ontology: tool for the unification of biology. The Gene Ontology Consortium. *Nat Genet* **25**, 25–29 (2000).

27. Gene Ontology Consortium. The Gene Ontology knowledgebase in 2026. *Nucleic Acids Res* **54**, D1779–D1792 (2026).

28. Kanehisa, M. & Goto, S. KEGG: kyoto encyclopedia of genes and genomes. *Nucleic Acids Res* **28**, 27–30 (2000).

29. Kanehisa, M., Furumichi, M., Sato, Y., Matsuura, Y. & Ishiguro-Watanabe, M. KEGG: biological systems database as a model of the real world. *Nucleic Acids Res* **53**, D672–D677 (2025).

30. Gargano, M. A. *et al.* The Human Phenotype Ontology in 2024: phenotypes around the world. *Nucleic Acids Res.* **52**, D1333–D1346 (2024).

31. Wamsley, B. *et al.* Molecular cascades and cell type-specific signatures in ASD revealed by single-cell genomics. *Science* **384**, eadh2602 (2024).

32. Wickham, H. *et al.* Welcome to the Tidyverse. *Journal of Open Source Software* **4**, 1686 (2019).

33. Wickham, H. *ggplot2: Elegant Graphics for Data Analysis*. (Springer, 2016).

34. Wickham, H., Pedersen, T. L. & Seidel, D. Scales: Scale functions for visualization. *CRAN: Contributed Packages* The R Foundation https://doi.org/10.32614/cran.package.scales (2011).

35. Szklarczyk, D. *et al.* The STRING database in 2023: protein-protein association networks and functional enrichment analyses for any sequenced genome of interest. *Nucleic Acids Res*

**51**, D638–D646 (2023).
